# Quantitative physiology and biomass composition of *Cyberlindnera jadinii* in ethanol-grown cultures

**DOI:** 10.1101/2024.08.31.610600

**Authors:** Marcel A. Vieira-Lara, Marieke Warmerdam, Erik A. F. de Hulster, Marcel van den Broek, Jean-Marc Daran, Jack T. Pronk

## Abstract

**Background:** Elimination of greenhouse gas emissions in industrial biotechnology requires replacement of carbohydrates by alternative carbon substrates, produced from CO_2_ and waste streams. Ethanol is already industrially produced from agricultural residues and waste gas and is miscible with water, self-sterilising and energy-dense. The yeast *C. jadinii* can grow on ethanol and has a history in the production of single-cell protein (SCP) for feed and food applications. To address a knowledge gap in quantitative physiology of *C. jadinii* during growth on ethanol, this study investigates growth kinetics, growth energetics, nutritional requirements, and biomass composition of *C. jadinii* strains in batch, chemostat and fed-batch cultures.

**Results:** In aerobic, ethanol-limited chemostat cultures, *C. jadinii* CBS 621 exhibited a maximum biomass yield on ethanol (Y_x/s_^max^) of 0.83 g_biomass_ (g_ethanol_)^-1^ and an estimated maintenance requirement for ATP (m_ATP_) of 2.7 mmol·(g_biomass_)^-1^·h^-1^. Even at specific growth rates below 0.05 h^-1^, a stable protein content of approximately 0.54 g_protein_·(g_biomass_)^-1^ was observed. At low specific growth rates, up to 17% of the proteome consisted of alcohol dehydrogenase proteins, followed by aldehyde dehydrogenases and acetyl-CoA synthetase. Of 13 *C. jadinii* strains evaluated, 11 displayed fast growth on ethanol (μ_max_ > 0.4 h^-1^) in mineral medium without vitamins, and CBS 621 was found to be a thiamine auxotroph. The prototrophic strain *C. jadinii* CBS 5947 was grown on an inorganic salts medium in fed-batch cultures (10-L scale) fed with pure ethanol. Biomass concentrations in these cultures increased up to 100 g_biomass_·(kg_broth_)^-1^, with a biomass yield of 0.65 g_biomass_·(g_ethanol_)^-1^. Model-based simulation, based on quantitative parameters determined in chemostat cultures, adequately predicted biomass production. A different protein content of chemostat- and fed-batch-grown biomass (54% and 42%, respectively) may reflect the more dynamic conditions in fed-batch cultures.

**Conclusions:** Analysis of ethanol-grown batch, chemostat and fed-batch cultures provided a quantitative physiology baseline for fundamental and applied research on *C. jadinii*. Its high maximum growth rate, high energetic efficiency of ethanol dissimilation, simple nutritional requirements and high protein content, make *C. jadinii* a highly interesting platform for production of SCP and other products from ethanol.

## Introduction

Establishing a circular, net-zero emission industrial biotechnology industry requires replacing refined sugars, which are currently the predominant carbon sources for industrial fermentation processes, with feedstocks derived from CO_2_ and renewable waste streams (1, 2). Especially for products that require ATP-intensive synthesis, achieving high titres, rates, and yields can be facilitated by first converting primary carbon sources into smaller organic molecules. These molecules can then serve as substrates in separate, intensive aerobic fermentation processes (1). In addition to microbial fermentation of lignocellulosic residues from agriculture and forestry (3), such small molecules can be produced by gas fermentation (4), electrochemical production from CO_2_ (1, 5) and, potentially, from microbial electrosynthesis (6, 7). The resulting organic molecules include C_1_ compounds such as formic acid and methanol as well as the C_2_ compounds acetic acid and ethanol (1).

Ethanol is an energy-dense substrate that is completely miscible in water and, at high concentrations, self-sterilising. Especially at low pH, ethanol is less toxic than other C_1_ and C_2_ compounds generated from CO_2_. Ethanol can be produced electrochemically from CO_2_ at lab-scale with a Faraday efficiency of 68% (8). Production of ethanol from agricultural residues by yeast (0.3-0.5 g_ethanol_·(g_sugar_)^-1^) (9) and from waste gases by *Clostridium autoethanogenum* (up to 200 mmol_ethanol_·(g_biomass_)^-1^ and 30% carbon recovery in ethanol) (4, 10, 11) have both transitioned to industrial scale. Ethanol can be readily assimilated by a variety of microorganisms (12), including yeast species that are already applied in large-scale sugar-based fermentation processes. However, compared to the wealth of information on quantitative physiology of sugar-grown aerobic cultures of industrially relevant yeast species, quantitative information on the stoichiometry, kinetics, and energetics of ethanol-grown cultures of such yeasts is scarce (13).

After diffusion over the yeast plasma membrane (14), ethanol is first converted to acetyl-CoA. This conversion involves the enzymes NAD^+^-dependent alcohol dehydrogenase, NAD(P)^+^-dependent acetaldehyde dehydrogenase and acetyl-CoA synthetase. Further dissimilation and assimilation of acetyl-CoA involves the tricarboxylic acid (TCA) cycle, glyoxylate cycle and gluconeogenesis. Dissimilation of ethanol to CO_2_ generates reduced cofactors (5 NAD(P)H and 1 FADH_2_) and requires input of one ATP-equivalent due to the combination of the investment of 2 ATP equivalents in the acetyl-CoA synthetase reaction (15) and the formation of 1 ATP equivalent by substrate-level phosphorylation in the TCA cycle. Energy coupling of the subsequent reoxidation of NADH and FADH_2_ by mitochondrial respiration has a strong impact on the net ATP yield from ethanol dissimilation and, thereby, on product yields on ethanol and on oxygen. The efficiency of this energy coupling is expressed by the P/O ratio, which indicates how many moles of ATP oxidative phosphorylation generates when a single electron pair (for example derived from NADH) is transferred to oxygen.

Involvement of an active multi-subunit proton-pumping NADH dehydrogenase (‘Complex I’) in the mitochondrial respiratory chains of yeasts strongly influences their P/O ratio (16, 17). Several industrially relevant yeasts, such as *Saccharomyces cerevisiae* and *Kluyveromyces lactis*, lack the genetic information for synthesizing Complex I and, instead, only harbour single-subunit, non-proton-translocating NADH dehydrogenases (18, 19). Other yeasts encode the genetic information for synthesizing both Complex I and non-proton-translocating NADH dehydrogenases (20). In such yeasts, energy coupling of respiration depends on how regulation of protein synthesis and/or activity determine the distribution of NADH oxidation over Complex I and less efficient systems.

The ascomycete budding yeast *Cyberlindnera jadinii* can grow on a variety of carbon sources, including sugars, organic acids and alcohols, and on nitrogen sources including ammonium, nitrate, urea and amino acids (21). Although no extensive quantitative physiology studies have been published on ethanol-grown cultures of *C. jadinii*, this yeast has been reported to show high biomass yields on ethanol (g_biomass_·(g_ethanol_)^-1^) (18). This observation is consistent with the presence, in the genome of *C. jadinii*, of the genetic information for encoding a Complex I NADH dehydrogenase (22). Based on dry weight, glucose-grown biomass of *C. jadinii* contains approximately 50% protein (18). *C. jadinii* strains have been reported to grow without vitamin supplementation (23), which enables simple processes and may offer options to select strains with high vitamin content.

*C. jadinii* is a polyploid teleomorph of *Candida utilis* (21, 24), which has a long history of safe use in biotechnology. *C. jadinii* biomass can be used as single-cell protein (SCP) as a food ingredient (25) and for animal nutrition (26). Nutritional studies have shown that *C. jadinii* SCP can functionally replace conventional protein sources in fish, chicken and swine feed (27–29). A historical example of SCP is Pruteen®, a product launched by Imperial Chemical Industries (ICI) in 1980, produced from hydrocarbon-derived methanol using *M. methylotrophus* (Matassa et al., 2016; Westlake, 1986). Although Pruteen® production was discontinued for economic reasons, interest in SCP as a viable protein source in animal feed continues to grow given the emergence of alternative feedstocks (30, 31). SCP has the potential to alleviate negative environmental impacts of food systems (32–35) due to a lower carbon footprint, water use and land usage than those of other protein sources (36).

Large-scale industrial production of microbial biomass typically involves aerobic fed-batch cultures. Because of oxygen transfer limitations in industrial bioreactors, achieving high biomass concentrations in such processes requires a decreasing specific growth rate profile (37). For understanding and predicting microbial performance in such industrial contexts, knowledge on growth-rate-dependent physiology, with special attention for maintenance-energy requirements, biomass yields on ethanol and oxygen, and, in the case of SCP production, biomass composition is essential. This study therefore aims to investigate the quantitative physiology of *C. jadinii* with ethanol as a substrate and its implications for SCP production. To this end, we studied growth stoichiometry, growth energetics, vitamin requirements and biomass composition in batch and chemostat cultures of *C. jadinii* strains. Based on quantitative information derived from these experiments, a fed-batch cultivation protocol for SCP production from pure ethanol as carbon source was designed and conducted at 10-L scale.

## Materials & Methods

### Yeast strains and media

*Cyberlindnera jadinii* strains CBS 621, CBS 567, CBS 841, CBS 842, CBS 890, CBS 1516, CBS 1517, CBS 1600, CBS 2160, CBS 4511, CBS 5609, CBS 5947, and CBS 7232 were obtained from the Westerdijk Institute (Utrecht, The Netherlands). *Saccharomyces cerevisiae* CEN.PK113-7D was provided by Dr. Peter Kötter (Goethe University, Frankfurt, Germany). Cultures were grown on a synthetic medium (SM) containing 5 g·L^-1^ (NH_4_)_2_SO_4_, 3 g·L^-1^ KH_2_PO_4_ 0.5 g·L^-1^ MgSO_4_·7H_2_O, 15 mg·L^-1^ EDTA, 4.5 mg·L^-1^ ZnSO_4_·7H_2_O, 0.3 mg·L^-1^ CoCl_2_·6H_2_O, 1 mg·L^-1^ MnCl_2_·H_2_O, 0.3 mg·L^-1^ CuSO_4_·5H_2_O, 4.5 mg·L^-1^ CaCl_2_·2H_2_O, 3 mg·L^-1^ FeSO_4_·7H_2_O, 0.4 mg·L^-1^ Na_2_MoO_4_·2H_2_O, 1 mg·L^-1^ H_3_BO_3_, 0.1 mg·L^-1^ KI. The pH of the medium was adjusted to 6.0 with 1 M KOH. After autoclaving at 120 °C for 20 min, 1 mL·L^-1^ of a filter-sterilized vitamin solution (0.05 g·L^-1^ D-(+)-biotin, 1.0 g·L^-1^ D-calcium pantothenate, 1.0 g·L^-1^ nicotinic acid, 25 g·L^-1^ *myo*-inositol, 1.0 g·L^-1^ thiamine hydrochloride, 1.0 g·L^-1^ pyridoxal hydrochloride, 0.2 g·L^-1^ 4-aminobenzoic acid) (38) was aseptically added. In some experiments, individual or all vitamins were omitted as indicated. Glucose solutions autoclaved at 110 °C for 20 min or pure ethanol were aseptically added to the medium along with the vitamin solution. For strain storage, 30% (v/v) glycerol (final concentration) was added to shake-flask cultures grown overnight at 200 rpm and 30 °C in YPD medium (10 g·L^-1^ yeast extract, 20 g·L^-1^ peptone and 20 g·L^-1^ glucose) and 1 mL samples were kept at -80 °C.

### Shake flask and microtiter plate cultivation

Shake-flask cultures, grown in 500-mL flasks containing 100 mL SM supplemented with either 20 g·L^-1^ D-glucose (SMD) or 7.5 g·L^-1^ ethanol (SME), were incubated at 30 °C and 200 rpm in Innova incubators (Brunswick Scientific Edison, NJ). Initial cultures were inoculated from frozen glycerol samples, incubated overnight, and used for a second inoculation in the same medium. Upon reaching early exponential phase, these pre-cultures were used to inoculate a third culture, either in shake flasks or microtiter plates, at an initial OD_660_ of 0.2-1.0. Absorbance at 660 nm (OD_660_) was measured with a Jenway 7200 spectrophotometer after accurate dilution of samples to yield OD_660_ values between 0.1 and 0.3. Specific growth rates were calculated by fitting exponential phase data (OD 0.2 – 5.0) to the equation *ln OD(t) = ln OD*(*0*) *+ µ_max_*·*t*, in which *µ_max_* (h^-1^) represents the maximum specific growth rate. Cultivation in 96-well plates was done in a Growth Profiler 960 (Enzyscreen BV, Heemstede, the Netherlands), operated at 30 °C and at 250 rpm. Images from the bottom of the incubator plates were taken every 20 min. The number of green pixels per picture (Green Value) was quantified with the GP960Viewer software. Green Values correlate with OD_660_ and were used to calculate maximum specific growth rates from serial dilutions (39). Mathematical details and assumptions of the method are described in Supp. Material S3.

### Chemostat cultivation

Chemostat cultures of *C. jadinii* and *S. cerevisiae* were grown at 30 °C in 1.5 L bioreactors connected to an ez-Control (Applikon Getinge, Delft, The Netherlands). Glucose- and ethanol-limited cultures were grown on SM with 5 g·L^-1^ glucose and 3.75 g·L^-1^ ethanol, respectively, unless otherwise stated, with 0.2 g·L^-1^ Pluronic PE 6100 anti-foaming agent (BASF, Ludwigshafen, Germany). The working volume of cultures was kept at 1.0 L with an electrical level sensor connected to the effluent pump. Unless stated otherwise, culture pH was maintained at 5.0 by automatic addition of 2 M KOH. Cultures were stirred at 800 rpm with 2 Rushton impellers and sparged with air at 0.5 L/min. CO_2_ and O_2_ contents in the in- and off--gas were measured with a multi-gas analyser (MultiExact 4100 Multi-gas Analyser, Servomex). Dissolved oxygen concentration was monitored with an autoclavable oxygen electrode (Applisens, Schiedam The Netherlands). After inoculation and a batch cultivation phase, the continuous cultivation mode was initiated when a sharp decrease of the CO_2_ output indicated substrate depletion. At least two pre-steady-state samples were taken from the effluent line when at least 5 volume changes had passed since the onset of continuous cultivation. When values of dry weight and CO_2_ differed by less than 5% for two consecutive volume changes, steady-state samples were collected directly from the culture. Carbon recoveries, calculated based on a yeast biomass carbon content of 45.5% (w/w) (40) were between 95 and 105% unless stated otherwise. Maintenance coefficient and maximum biomass yield on substrate were calculated by linear regression of substrate consumption rates (*q_s_*) plotted against the dilution rate (D) according to the Pirt equation (eq. 1; (41).

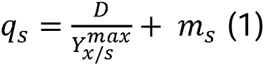

### Fed-batch cultivation

Fed-batch cultivation was conducted at 30 °C in 20-L Bio Bench reactors (Applikon Getinge, Delft, The Netherlands) with an initial working volume of 8.0 L. Stirrer speed was kept at 800 rpm and pressurised air (20.95% O_2_, 0.04% CO_2_) was supplied at a flow rate of 7 L·min^-1^. An overpressure of 1.0 bar (2.0 bar total pressure) was applied throughout cultivation to enhance oxygen transfer. A polarographic oxygen electrode (model 32 275 6800; Ingold) was used to monitor the dissolved oxygen concentration to ensure it remained above 15% of air saturation (42).

The medium used for fed-batch cultivation contained 2 g·L^-1^ (NH_4_)_2_SO_4_, 20 g·L^-1^ KH_2_PO_4_ 6 g·L^-1^ MgSO_4_·7H_2_O, and trace elements (150 mg·L^-1^ EDTA, 45 mg·L^-1^ ZnSO_4_·7H_2_O, 3 mg·L^-1^ CoCl_2_·6H_2_O, 10 mg·L^-1^ MnCl_2_·H_2_O, 3 mg·L^-1^ CuSO_4_·5H_2_O, 45 mg·L^-1^ CaCl_2_·2H_2_O, 30 mg·L^-1^ FeSO_4_·7H_2_O, 4 mg·L^-1^ Na_2_MoO_4_·2H_2_O, 10 mg·L^-1^ H_3_BO_3_, 1 mg·L^-1^ KI) and 0.2 g·L^-1^ Pluronic PE 6100 antifoam. Ammonium hydroxide (25% w/v) was used to control culture pH at 5.0 and to simultaneously provide on-demand nitrogen-source supply. For the initial batch phase, which was inoculated at an OD_660_ of 0.2, 12 g·L^-1^ ethanol was used as carbon source. When the end of the exponential growth phase was indicated by a simultaneous decrease of CO_2_ production and an increase in the dissolved-oxygen concentration, the fed-batch phase was initiated. During this phase, absolute ethanol was used as carbon source and pumped into the reactor with a controllable Masterflex model 7518-00 pump (Cole-Parmer Instrument Company, Chicago, IL). The feed rate was controlled by a MFCS SCADA Software (Sartorius Stedim Biotech GmbH, Göttingen, Germany) according to equation 2. Over the duration of the fed batch phase, this feed profile resulted in flow rates between 0.006 and 0.050 kg_ethanol_·h^-1^.

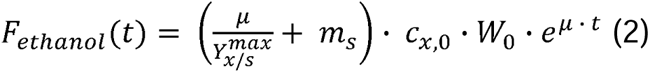

In equation 2, *F_ethanol_* indicates the feed rate of ethanol in kg_ethanol_·h^-1^, c_x,0_ the initial biomass concentration (g_biomass_·(kg_broth_^-1^)), W_0_ the initial culture weight (kg_broth_) and t the time (h) after starting the feed. 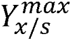 is the maximum biomass yield on substrate in the absence of maintenance (g_biomass_·(g_ethanol_^-1^)) and *m_s_*is the maintenance coefficient (g_ethanol_·(g_biomass_^-1^)·h^-1^), both calculated from chemostat experiments. The specific growth rate was kept at 0.10 h^-1^ until the dissolved oxygen concentration decreased to 15% of air saturation. At that point, the feed was changed from exponentially increasing to constant. The fed-batch process was simulated in Python essentially as described by Hensing et al., 1995 (37). The volumetric mass transfer coefficient (*k_L_a*) (43) was experimentally determined to be 210 ± 11 h^-1^ with the dynamic technique of absorption (44), and a value of 200 h^-1^ was used for the simulation. Scripts are found in Appendix 1.

### Analytical methods

Biomass dry weight in culture samples was measured as previously described (45). Briefly, dried nitrocellulose filters (pore size 0.45 µm, Gelman Laboratory, Ann Arbor, MI) stored at 80 °C were weighed and used to filter 10.0 mL culture samples. Fed-batch samples were diluted in demineralized water to a concentration < 5 g_biomass_ · (kg_broth_)^-1^ prior to dry weight measurements. To correct for increased broth density at the end of the process, density was measured by weighing 10.0 mL of broth and used to convert values from g_biomass_ · (L_broth_)^-1^ to g_biomass_· (kg_broth_)^-1^ (for samples with > 70 g_biomass_ · (kg_broth_)^-1^). After washing with demineralized water, filters were dried for 20 min at 350 W output in a microwave (Bosch, Stuttgart, Germany) and weighed again. For determination of metabolite concentrations in culture supernatants, samples were rapidly quenched with cold steel beads and filtered as described by Mashego et al., 2003 (46). Concentrations of glucose and metabolites were determined by HPLC (47). Biomass-specific rates of production and consumption of substrates and products, respectively, as well as carbon recoveries, were calculated with a script available from Appendix 2.

### Determination of biomass composition

Protein content of yeast biomass was determined with the Biuret method according to Verduyn et al. (16), and lipid content according to Izard & Limberger, and Johnson et al. (48, 49) with modifications. Briefly, cell suspension (20 mL) was centrifuged at 10,000 g for 5 min. The pellet was washed and resuspended in demineralized water to give a total volume of 10.0 mL H_2_O and stored at -20 °C. For protein measurements, thawed samples were mixed with 3 M NaOH (2:1), boiled at 100 °C for 10 min and cooled on ice. After adding CuSO_4_·5H_2_O (2.5% w/v) to samples (1:3), they were incubated at room temperature for 5 min and centrifuged at 13,000 rpm for 5 min. The absorbance of the clear supernatant was then measured at 510 nm. A solution of 5 g·L^-1^ bovine serum albumin (Merck-Sigma A9647) was used to construct a standard curve. For total lipid determination, 100 µL of sample were mixed with 1.0 mL H_2_SO_4_ and vortexed. Samples were incubated at 100 °C for 10 min and cooled for 5 min. Subsequently, 2.5 mL of phosphoric acid-vanillin reagent (1.2 g·L^-1^ vanillin and 68% phosphoric acid) were added, samples were vortexed and incubated at 37 °C for 15 min. Samples were cooled to room temperature and absorbance was read at 530 nm. Standard solutions of sunflower oil (Merck-Sigma S5007) in chloroform (0 – 1.0 mg/mL) were used to construct a calibration curve. Chloroform was fully evaporated prior to H_2_SO_4_ addition.

To measure RNA and carbohydrate contents of yeast biomass, culture samples were washed once with demineralized water and pellets containing 3 to 5 mg biomass were stored at -20 °C. A 2.0 g·L^-1^ solution of yeast RNA (Sigma 10109223001) was prepared in 10 mM Tris-HCl, pH 8.0 and also stored at -20 °C. Before analysis, the stock solution was diluted to obtain standard solutions with RNA concentrations ranging from 5 to 60 mg·L^-1^. RNA was extracted (50) and quantified by measuring absorbance at 260 nm (U-3010 spectrophotometer, Hitachi). Total carbohydrate contents in biomass were quantified according to (51). Briefly, 5 mL of concentrated H_2_SO_4_ were added to 1 mL of sample, followed by the addition of 1 mL of phenol solution (50 g·L^-1^). Samples were incubated at 100 °C for 15 min and subsequently cooled to room temperature. Absorbance was measured at 488 nm. A standard solution of glucose and mannose (2:1) was used to construct a calibration range (0.0 - 0.2 g·L^-1^). To correct for the presence of nucleic acid pentoses, relative extinction coefficients of 0.36 and 0.26 (versus the coefficient obtained from the standard curve) were used for RNA and DNA, respectively (adapted from (40)). The following formula was used:

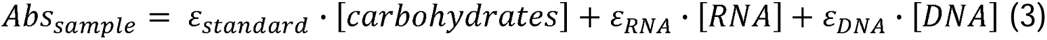

Where *Abs* stands for absorbance, ε_standard_ was obtained from freshly prepared standard curves, RNA content was measured, and DNA content was assumed to be 0.5% of the biomass dry weight for all samples.

### Proteome analysis

Biomass samples (20 mL, containing approximately 3 g biomass·L^-1^) were centrifuged and washed once with demineralized water. Pellets were resuspended in cold methanol (-20°C) and lysed with a Precellys Homogenizer (Bertin technologies SAS, Montigny-le-Bretonneux, France) at 6,800 rpm for 15 s. Total protein concentrations of the resulting samples were measured by Qubit protein assay, and proteins were isolated by Bligh and Dyer extraction. Isolated proteins were processed further by reduction, alkylation, and trypsin digestion (52). Samples were analysed in technical triplicates by liquid chromatography coupled to tandem mass spectrometry (LC-MS/MS) using a Vanquish UHPLC coupled to a Qexactive Plus (Thermo Fisher Scientific, Waltham, USA). Peptides were separated using reverse-phase chromatography with a gradient of water with 0.1% formic acid (solvent A) and 20% water and 0.1% formic acid in acetonitrile (solvent B) from 5% B to 60% B in 52 min. Data-independent acquisition (DIA) was performed with a resolution setting of 17,500 within the 400 to 1,200 m/z range and a maximum injection time of 20 ms, followed by high-energy collision-induced dissociation activated (HCD) MS/MS scans with a resolution setting of 17,500 and injection time set to auto. Four DIA MS/MS fragmentation scans were performed after each full scan and the mass range was divided into 100 m/z windows. Raw files were analysed with the Pulsar search engine and Spectronaut, version 17.5, against the annotated proteins of *C. jadinii* (UniProtKB, strain ATCC 18201/CBS 1600 [www.uniprot.org]). Label-free quantification was performed using the top three peptides measured for each protein. Retention time realignment was based on local (non-linear) regression.

The obtained signal per protein (*raw signal (i)*, proportional to protein copy number) was multiplied by the respective molecular weight, yielding a *signal (i)*. Subsequently, the *signal (i)* obtained per protein was divided by the sum of all *signals (i)*’s obtained in the sample and multiplied by 100. This resulted in the approximate percentage (in weight/weight) by which each protein contributed to the total proteome mass.

### Amino acid analysis

Culture samples were washed once with demineralized water and pellets containing approximately 50 mg biomass were stored at -20 °C. Total amino acid content of yeast biomass was determined after hydrolysis of the biomass according to the Accq-Tag method (Waters). Before hydrolysis, samples were weighed in a headspace vial, which was subsequently flushed with nitrogen to remove oxygen. Hydrolysis with 6 M HCl was performed to release free amino acids from proteins and peptides. Free amino acids were derivatized on the NH_2_ terminus with (AQC) 6-aminoquinolyl-N-hydroxysuccinimidyl carbamate. For complete conversion, samples were incubated for 10 min at 55 °C. Derivatized amino acids were analysed using UPLC reversed-phase chromatography with diode array detection at 260 nm (Acquity UPLC, Waters Corporation, Milford, USA). Derivatized amino acids were chromatographically separated with an Acquity UPLC BEH C18 1.7 µm 2.1×100 mm (Waters) analytical column and a gradient with Mobile phase A: AccQ·Tag Ultra Eluent A, and Mobile phase B: AccQ·Tag Ultra Eluent B, starting at 0.1% B to 9.1% B in 5.74 min increasing to 21.2% B at 7.74 min followed by a ramp to 59.6% B at 8.04 min with a flow rate of 0.7 mL/min. Glutamine, asparagine, tryptophan and cysteine could not be determined with this method because glutamine and asparagine are converted to glutamic acid and aspartic acid, and tryptophan breaks down during hydrolysis.

### Whole-genome sequencing

Genomic DNA was isolated from culture samples containing approximately 7.0·10^9^ cells with the Qiagen 100/G kit (Qiagen, Hilden, Germany), according to manufacturer’s instructions. Genomic DNA of *C. jadinii* CBS 621 was sequenced in-house on an Illumina MiSeq sequencer (Illumina, San Diego, CA) to obtain a paired-end library with an insert-size of 550 bp and 300 bp read length using TruSeq Nano DNA library preparation, yielding 6.4 Gb of genomic DNA sequence. *C. jadinii* CBS 5947 was sequenced at Macrogen on an Illumina NovaSeq sequencer to obtain a paired-end library with an insert-size of 550 bp and 151 bp read length using TruSeq DNA PCR-free library preparation, yielding 3.5 Gb of genomic DNA sequence. Reads of strains CBS 621 and CBS 5947 were mapped against the *C. jadinii* NBRC 0988 genome (BioProject: PRJDB11630; GCA_024346625.1) using BWA (version 0.7.15) (53). Alignments were processed using SAMtools (version 1.3.1.) (54) and variants were called by applying Pilon (version 1.18) (55). Heterozygous single-nucleotide polymorphism (SNP) ratios of each chromosome were extracted from the Pilon produced variant calling file (VCF) and the distributions were plotted by using ggplot2 (56). *De novo* assembly was performed on the Illumina reads with the SPAdes genome assembler (version 3.9.0) (57). Sequencing data are available at NCBI (https://www.ncbi.nlm.nih.gov/) under the BioProject accession number PRJNA1153750.

### Vitamin measurements

Analyses of selected vitamins (thiamine, pyridoxine, nicotinic acid and riboflavin) were conducted via Elisa by SGS Nederland B.V., division SGS Food Lab. Biomass samples containing circa 3 g_biomass_ were used.

### Software and Statistics

Scripts used for calculations were written in Python (v3.10), using Jupyter Notebook (v 6.4.11). Data visualisation and statistical analysis were conducted in GraphPad Prism (v9.0, GraphPad Software, San Diego, CA).

## Results

### 1. Biomass yield and protein content in glucose- and ethanol-grown chemostat cultures

To verify earlier reports that *C. jadinii* shows higher biomass yields on ethanol than *S. cerevisiae* (16, 18), aerobic ethanol-limited chemostat cultures of the model strains *C. jadinii* CBS 621 (18, 58, 59) and *S. cerevisiae* CEN.PK113-7D (40, 60–62) were grown at a dilution rate of 0.10 h^-1^. Consistent with earlier reports (18), steady-state cultures of *S. cerevisiae* CEN.PK113-7D showed a biomass yield of 0.59 ± 0.01 g_biomass_·(g_ethanol_)^-1^ (Fig 1A). In four independent chemostat cultures of *C. jadinii* CBS 621, biomass yields were consistently 23% higher (0.74 ± 0.01 g_biomass_·(g_ethanol_)^-1^) than in those of *S. cerevisiae*. The biomass yields on ethanol of *C. jadinii* CBS 621 were higher than previously reported for chemostat cultures of this strain (0.69 g_biomass_·(g_ethanol_)^-1^) (16, 18). In contrast to the ethanol-limited cultures, glucose-limited chemostat cultures of *C. jadinii* CBS 621 and *S. cerevisiae* CEN.PK113-7D showed only 6% difference in biomass yield (0.51 ± 0.01 g_biomass_·(g_glucose_)^-1^ and 0.48 ± 0.00 g_biomass_·(g_glucose_)^-1^, respectively; Fig 1A). Both species showed lower biomass yields on oxygen in ethanol-limited cultures than in glucose-limited cultures (Fig 1B). However, biomass yields on oxygen in ethanol-limited cultures of *C. jadinii* were approximately 50% higher in *C. jadinii* than *S. cerevisiae* (Table S1). The different biomass yields of *C. jadinii* and *S. cerevisiae* on ethanol may reflect the presence and absence, respectively, of a proton-pumping respiratory complex I in these two yeast species (16, 22).

**Figure 1:**
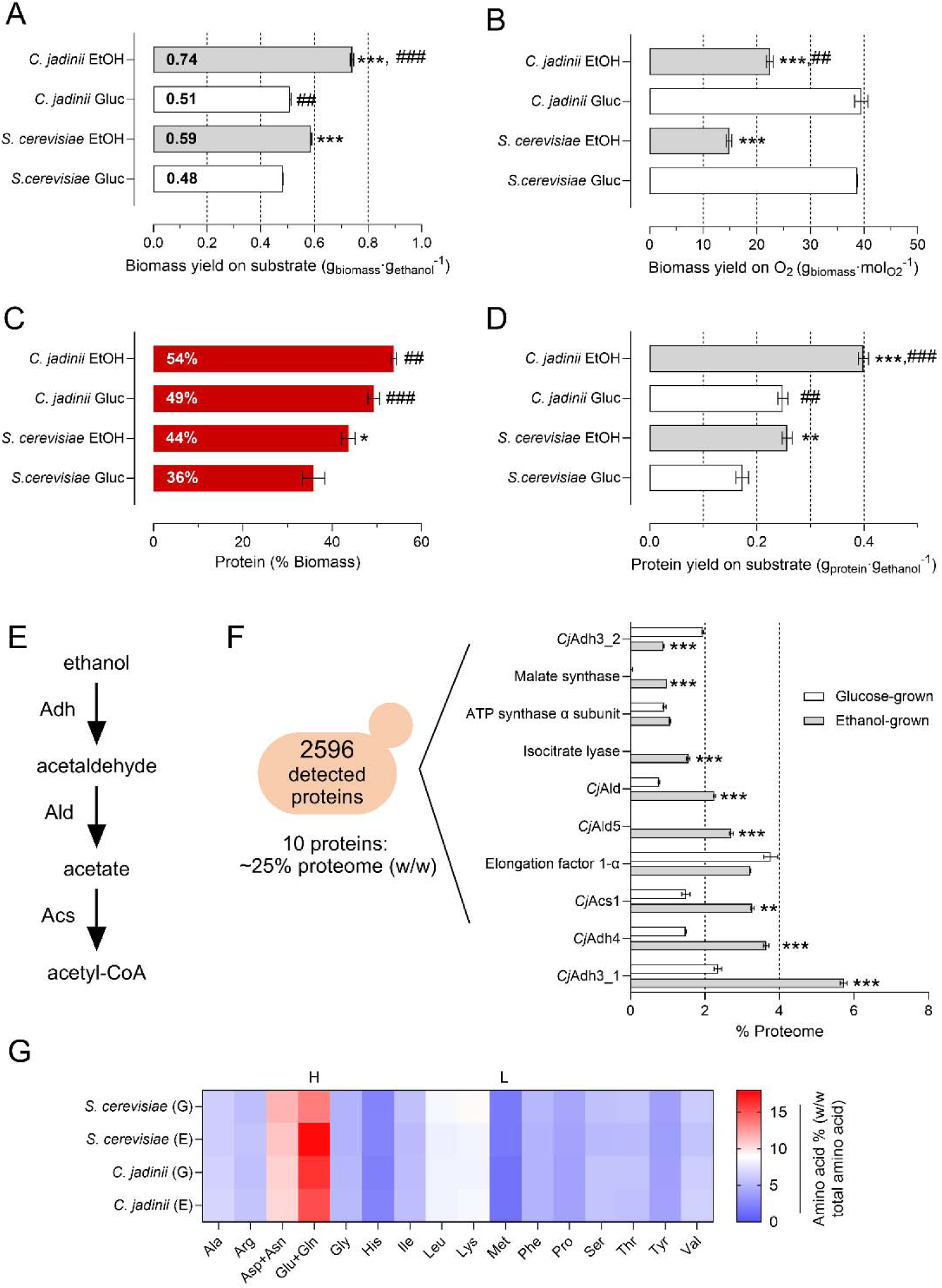
Effect of growth-limiting carbon source on the physiology of *C. jadinii* and proteome composition. *S. cerevisiae* CEN.PK113-7D and *C. jadinii* CBS 621 were grown in steady-state, aerobic ethanol-limited and glucose-limited chemostats at a dilution rate of 0.10 h^-1^. (A, B) Biomass yields on substrate and oxygen, respectively. (C) Protein content as a percentage of biomass. (D) Protein yields on substrate. (E) Enzymes involved in conversion of ethanol to acetyl-CoA in yeasts. Adh: alcohol dehydrogenase, Ald: aldehyde dehydrogenase, Acs: acetyl-CoA synthetase. (F) Top 10 most abundant proteins (with at least 3 identified peptides) in the proteome of ethanol-grown *C. jadinii* versus the levels of the same proteins in glucose-limited chemostats in the same species. *Cj* stands for *Cyberlindnera jadinii*. (G) Protein amino acid residue composition (as a percentage of the total amino acid residue pool). H stands for highest, and L for lowest amino acids in abundance for all analysed species and conditions. #p<0.05, ##p<0.01, ###p<0.001, *S. cerevisiae* vs *C. jadinii*, same substrate. *p<0.05, **p<0.01, ***p<0.001, same yeast species, ethanol-(EtOH) vs glucose-grown (Gluc), n=2.

Consistent with earlier reports (18, 40), the highest protein content in yeast biomass (0.54 g_protein_·(g_biomass_)^-1^) was observed in the ethanol-limited *C. jadinii* cultures and the lowest protein content in the glucose-limited *S. cerevisiae* cultures (0.36 g_protein_·(g_biomass_)^-1^) (Fig 1C). In the ethanol-limited cultures of *C. jadinii*, the protein yield on ethanol reached 0.40 g_protein_·(g_ethanol_)^-1^ (Fig 1D).

### 2. Proteome and amino-acid composition of ethanol-grown *C. jadinii*

To further investigate the biomass composition of the ethanol- and glucose-limited chemostat cultures of *C. jadinii*, proteome analyses were performed. Of 2596 detected proteins, 1656 were identified with at least 3 different peptides (Supp. Material S1). In the ethanol-limited *C. jadinii* cultures, the 10 most abundant proteins comprised approximately 25% of the proteome. Two isoforms each of alcohol dehydrogenase (*Cj*Adh) and acetaldehyde dehydrogenase (*Cj*Ald), as well as acetyl-CoA synthetase 1 (*Cj*Acs1), which together catalyse the first three steps in ethanol catabolism (Fig 1E), accounted for 6 of the 10 most highly abundant proteins (Fig 1F). Key proteins in ethanol assimilation (isocitrate lyase and malate synthase) were also very highly expressed. While their levels were clearly higher in ethanol-grown cultures, the 6 proteins involved in ethanol catabolism were also abundant in cells grown under glucose-limitation (Fig 1F). In contrast, levels of elongation factor 1-α and the α-subunit of the F_1_F_o_-ATP synthase were similar in glucose-limited and ethanol-limited cultures (Fig 1F).

In view of the 75% higher respiration rate in ethanol-limited cultures than in glucose-limited cultures of *C. jadinii* CBS 621 (4.4 vs 2.5 mmol O_2_·(g_biomass_)^-1^·h^-1^, Table S1), levels of proteins involved in oxidative phosphorylation were also investigated. A set of 71 proteins involved in oxidative phosphorylation that were represented by at least 2 independent peptides and included subunits of all 4 respiratory complexes and F_1_F_o_-ATP synthase (Supp. Material S1) showed, on average, a 60% higher level in ethanol-limited cultures than in glucose-limited cultures. Complex I subunits (28 detected) were, on average, 75% higher in ethanol-compared to glucose-limited samples (Fig. S1A).

The amino acid composition of *C. jadinii*, which is important for food and feed applications of microbial biomass (31), was analysed and compared to that of *S. cerevisiae*. Biomass from ethanol- and glucose-grown chemostat cultures of the two yeasts showed similar amino acid composition, with high glutamate/glutamine levels (Table S2). A low content of methionine in glucose- and ethanol-grown chemostat cultures of both yeasts (1.35 to 1.80% of the proteome, Fig 1G) was previously reported for multiple yeasts species (27). The sum of the analysed amino acid contents followed the same trend as total protein measurements (Fig S1B) but resulted in lower absolute values (Fig S1C). There is a growing interest in using yeast biomass in aquaculture to produce fish feed and thereby to curtail overfishing for fishmeal (27, 31, 32). A comparison of amino acid levels in ethanol-grown *C. jadinii* biomass with reference values for essential amino acids in fish feed identified methionine as limiting (Fig S1D), with all other amino acids at or above required levels, assuming yeast biomass as the only diet component.

### 3. Maximum biomass yield and maintenance-energy requirement of *C. jadinii* in ethanol-limited chemostat cultures

To analyse how specific growth rate affects biomass yield and to estimate the maintenance energy requirements of *C. jadinii* CBS 621, ethanol-limited chemostat cultures were grown at different dilution rates. To define a range of dilution rates, maximum specific growth rates (µ_max_) of *C. jadinii* were determined in batch cultures grown on synthetic medium with ethanol or glucose as sole carbon source (Fig 2A). Values of µ_max_ on ethanol and glucose were 0.43 ± 0.01 h^-1^ and 0.57 ± 0.01 h^-1^, respectively (Fig 2B). Both values are higher than the reported µ_max_ of *S. cerevisiae* CEN.PK113-7D on synthetic medium with glucose (0.39 h^-1^) (63).

**Figure 2:**
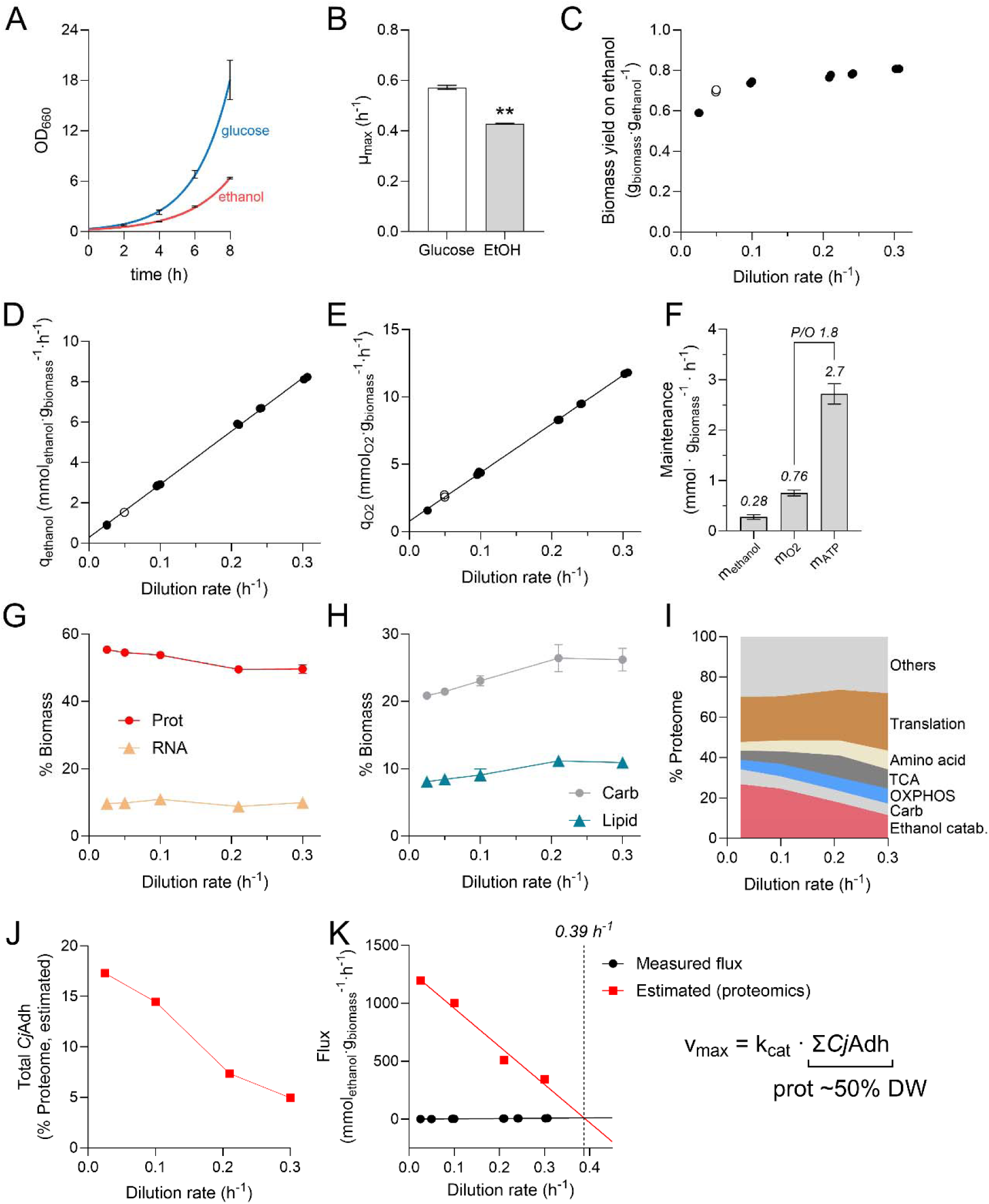
Growth rates and physiological characterization of *C. jadinii* in batch and ethanol-limited chemostat cultures at varying dilution rates. (A) Optical density at 660 nm (OD_660_) as a function of time and (B) maximum growth rates (µ_max_) of glucose- and ethanol-grown batch cultures. (C) Biomass yield on ethanol, (D) biomass-specific ethanol uptake rate (q_ethanol_) and (E) biomass-specific oxygen uptake rates (q_O2_) in aerobic, steady-state, ethanol-limited chemostat cultures of *C. jadinii* CBS 621. (F) Maintenance requirements for ethanol (m_ethanol_) and oxygen (m_O2_) calculated as the y-intercept of the fitted curves shown in D and E, respectively. Maintenance ATP requirements (m_ATP_) were estimated from m_O2,_ assuming an *in vivo* P/O ratio of 1.8 (16). (G) and (H) Quantification of the main components of the biomass in % of dry weight: proteins (Prot), RNA, carbohydrates (Carb) and lipids. (I) Protein allocation in ethanol-limited chemostat cultures grown at different dilution rates. Categories were manually curated and represent (in ascending order) Ethanol catabolism (first three steps shown in Fig 1F), Carbohydrate metabolism (Carb.), Oxidative Phosphorylation (OXPHOS), TCA cycle (including the glyoxylate shunt reactions), Amino acid metabolism, Translation and Others. The last category encompasses proteins from pathways with a lower proteome allocation or for which no pathway was assigned. (J) Total alcohol dehydrogenase protein (*Cj*Adh, % of proteome) as a function of dilution rate. (K) Estimated v_max_ for *Cj*Adh, assuming a k_cat_ of 150 min^-1^ for all isoforms (Brenda database (77), EC 1.1.1.1, average reported value for yeasts) and a proteome comprising 50% of the dry weight (w/w), plotted together with the ethanol consumption rates (q_ethanol_) from bioreactor experiments. 0.39 h^-1^ represents the intersection between both lines. Open symbols show carbon recovery of around 107%, while closed symbols represent carbon recoveries of 100±5%. **p<0.001, n=2.

Ethanol-limited chemostat cultures of *C. jadinii* were grown at dilution rates ranging from 0.025 to 0.30 h^-1^. Over this range of specific growth rates, biomass yields on ethanol ranged from 0.59 to 0.81 g_biomass_·(g_ethanol_)^-1^ (Fig 2C, Table S3). Using the Pirt equation (eq. 1), we derived maximum biomass yields on ethanol and oxygen based on the linear fits shown in Fig 2D-E. From the same data, maximum biomass yields of 0.83 g_biomass_·(g_ethanol_)^-1^ and 28 g_biomass_·(mol O_2_)^-1^ on oxygen were determined. The maximum yield on ethanol coincided with a theoretical value of 0.83 - 0.84 g_biomass_·(g_ethanol_)^-1^, estimated based on the assimilation equation for ethanol by *C. jadinii* (13, 16). This observation suggests that, at high specific growth rates, growth of *C. jadinii* CBS 621 is close to being carbon-limited rather than energy-limited. This means that the amount of ATP generated by respiratory reoxidation of the reduced redox cofactors formed during assimilation of ethanol almost exactly matches the amount of ATP needed for this assimilation process.

From Fig 2D-E, we derived maintenance coefficients for ethanol (m_s_) and oxygen (m_O2_) of 0.28 mmol_ethanol_·(g_biomass_)^-1^·h^-1^ and 0.76 mmol O_2_·(g_biomass_)^-1^·h^-1^, respectively (Fig 2F). Maintenance relates to use of energy substrate for processes that are not associated with growth, such as homeostasis of membrane potential and turnover of cellular components (64) and explains the decline in biomass yield as the dilution rate approaches zero (Fig 2C). Assuming an *in vivo* P/O ratio of 1.8 *for C. jadinii* (16) and growth-rate independent maintenance-energy requirements ((41), Eq. 1), we estimated an ATP requirement for maintenance (m_ATP_) of 2.7 mmol ATP·(g biomass)^-1^·h^-1^ (Fig 2F). This value is over 4-fold higher than the estimated m_ATP_ for *S. cerevisiae* grown aerobically in glucose-limited cultures (0.63 mmol_ATP_·(g_biomass_^-1^)·h^-1^) (60).

A previous publication reported that *C. jadinii* shows higher biomass yields at pH 4.0 than at pH 5.0 (65). However, we did not observe a meaningful change in biomass yield when ethanol-limited cultures were grown at pH 4.0 instead of 5.0, while protein contents of cultures grown at pH 4.0 were lower (Fig S2A-C). Co-feeding with formic acid, whose metabolism in yeasts is strictly dissimilatory, can supply additional NADH and thereby increase ATP availability for assimilation of the carbon source (18, 66, 67). Formic acid co-feeding was conducted in cultures grown at pH 5.0 (no expected toxicity (68)). This only marginally increased the biomass yield and protein content in ethanol-limited cultures, resulting in a protein yield on ethanol of 0.43 g _protein_·(g_ethanol_)^-1^ (Fig S2D-F). The effect of formic acid co-feeding was more pronounced in glucose-grown chemostat cultures, in which protein yields on glucose increased from 0.25 to 0.33 g_protein_·(g_glucose_)^-1^ (Fig S2F-H).

### 4. Impact of specific growth rate on protein content and alcohol dehydrogenase levels in ethanol-limited chemostat cultures

In aerobic, glucose-limited chemostat cultures of yeasts, the protein content of the biomass typically increases with increasing specific growth rate (40, 65, 69, 70). In contrast, aerobic ethanol-limited chemostat cultures of *C. jadinii* CBS 621 showed slightly lower protein contents with increasing growth rates (Fig 2G). Similarly, while RNA content was reported to positively correlate with specific growth rate in *S. cerevisiae* (40, 71), ethanol-limited chemostat cultures of *C. jadinii* CBS 621 showed essentially stable RNA contents across a range of dilution rates (Fig 2G). In contrast to protein and RNA contents, contents of carbohydrates and lipids in biomass increased with specific growth rate, representing 26 and 11%, respectively, of the biomass at D = 0.30 h^-1^ (Fig. 2H). Yields of all four macromolecular biomass components on ethanol showed a growth-rate dependent increase (Fig S2G-H), reflecting the trend of total biomass yields (Fig 2C).

Many studies report that the fraction of microbial proteomes allocated to the translation machinery increases with increasing specific growth rate (72–74). To investigate whether ethanol-grown *C. jadinii* exhibited a similar correlation, proteomics analyses were performed on samples from cultures grown at 0.025, 0.10, 0.21 and 0.30 h^-1^. The resulting proteome dataset (Supp. Material S2) was manually curated into 7 categories based on the Gene Ontology annotation obtained from UniProt (75). Levels of 357 proteins correlated well with specific growth rate (R^2^ > 0.90; 273 positive and 84 negative correlations; Supp. Material S2). The most prominent effect was a negative correlation of the abundance of proteins involved in ethanol catabolism (*Cj*Adh, *Cj*Ald and *Cj*Acs) with growth rate. These proteins occupied approximately 27% of the proteome at 0.025 h^-1^ and 12% at 0.3 h^-1^. In contrast, the fraction of proteins involved in oxidative phosphorylation (OXPHOS) increased from 4.8 to 7.4% (Fig. 2I), which positively correlated with oxygen consumption rates (Fig S3A). These abundances are similar to the proteome contribution of OXPHOS proteins of approximately 5% in *S. cerevisiae* and *Issatchenkia orientalis* (*Pichia kudriavzevii*) (72, 76). The fractions of TCA-cycle enzymes and proteins involved in amino acid synthesis both increased from approximately 4% at 0.025 h^-1^ to 9% at 0.3 h^-1^, while the fraction of the proteome allocated to the translation machinery increased from 22 to 29% (Fig 2I).

Of the proteins involved in ethanol catabolism, *Cj*Adh had the strongest impact on the composition of the proteome. Total *Cj*Adh levels (3 most abundant isoforms, Fig S3B) accounted for 17% and 5% of the proteome at 0.025 h^-1^ and 0.3 h^-1^, respectively (Fig 2J), followed by *Cj*Ald and *Cj*Acs (Fig S3C-D). Given the almost linear correlation between *Cj*Adh levels and dilution rate, we estimated *Cj*Adh’s specific maximum activity (v_max_). When these estimates were plotted against measured biomass-specific ethanol consumption rates (which should equal the *in vivo* flux through *Cj*Adh), the curves intercepted at D = 0.39 h^-1^ (Fig 2K), which is very close to the maximum specific growth rate of *C. jadinii* on ethanol (0.43 h^-1^). This comparison suggests that the maximum specific growth rate of *C. jadinii* CBS 621 on ethanol may be constrained at the proteome by the level of alcohol dehydrogenase.

### 5. *C. jadinii* CBS 621 is a thiamine auxotroph

The chemically defined medium used in the experiments described above contained seven B-type vitamins that are commonly included in defined media for yeast cultivation (78). However, although vitamin-independent growth of other *C. jadinii* strains has been reported (23), the vitamin dependence of strain CBS 621 has not been investigated. To assess vitamin requirements of this strain, it was grown in shake-flask cultures on ‘dropout’ synthetic media from which a single vitamin was omitted. In addition, growth was measured on a synthetic medium from which all 7 vitamins were omitted (Δvitamins). In these experiments, only the Δthiamine and Δvitamins media showed severely compromised growth (Fig 3A-B). When, after 24 h incubation in vitamin-free medium with ethanol, culture samples were used to inoculate new flasks, the growth deficiency after 24 h further incubation in vitamin-free medium was even more pronounced (Fig 3B). Addition of thiamine sufficed to restore growth (Fig 3C).

**Figure 3:**
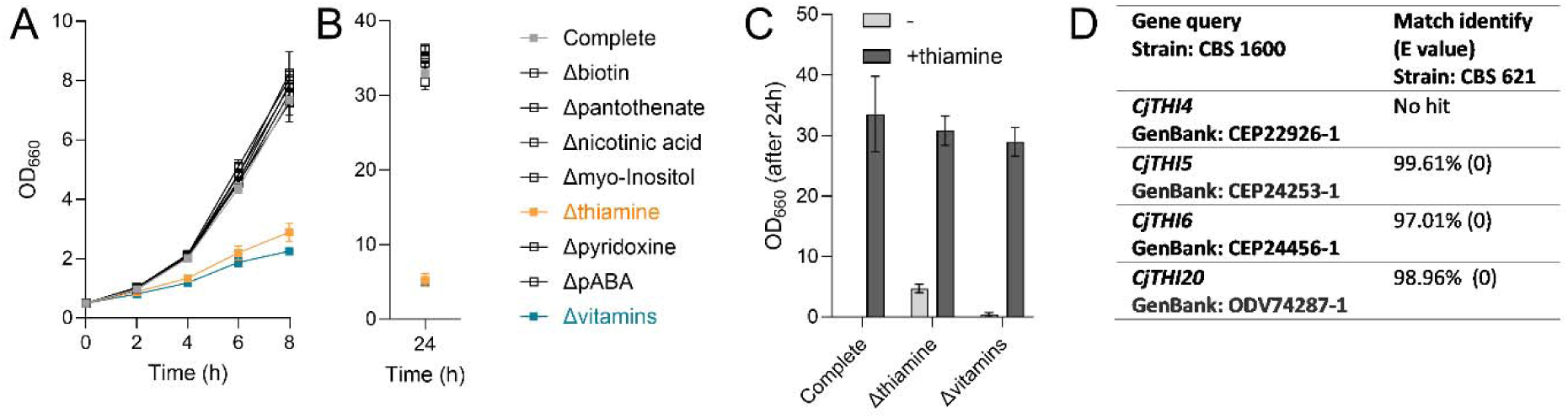
Growth of *C. jadinii* CBS 621 on single-vitamin dropout and vitamin-free media. (A) Growth curves of *C. jadinii* CBS 621 on single-vitamin dropout media in shake flasks grown on syntheti medium with ethanol as a sole carbon source. Δ“vitamin name” indicates absence of only the indicated vitamin from the standard vitamin mix. *Complete* indicates presence of all vitamins and Δ*vitamins* indicates absence of all vitamins. (B) OD_660_ after 24h of cultivation in shake flasks. (C) OD_660_ after further 24h of cultivation in shake flasks for samples from indicated groups (x-axis) that were reinoculated at OD 0.5 in the presence or absence of thiamine, n=2. (D) Overview of tBLASTn search to identify genes coding for enzymes in the biosynthesis of thiamine in CBS 621. The reference GenBank IDs used as queries were retrieved from UniProt, based on searches containing the respective protein name and species (*C. jadinii* CBS 1600). The tBLASTn algorithm was run locally using default settings.

To investigate the apparent thiamine auxotrophy *C. jadinii* CBS 621, its genome was sequenced and *de novo* assembled. The estimated genome size of 17.9 Mb (Table S4) is larger than that of other *C. jadinii* strains (21). This may be due to its high degree of heterozygosity, which generated alternative alleles for heterozygous regions during assembly. Based on the distribution of SNPs in the heterozygous regions across the genome, CBS 621 is most probably triploid (Fig S4A). A search for genes encoding proteins in thiamine biosynthesis (*CjTHI4*, *CjTHI5*, *CjTHI6* and *CjTHI20*) (78) was performed in UniProt, using *C. jadinii* as a taxonomy identifier. The annotated amino acid sequences for the type strain CBS 1600 were then used to identify homologs in strain CBS 621 with the BLAST tool (Fig 3D). This homology search yielded matches in strain CBS 621 with *CjTHI5*, *CjTHI6* and *CjTHI20*, but not with *CjTHI4*, which encodes thiamine thiazole synthase, a single-turnover ‘suicide enzyme’ (78) involved in thiamine biosynthesis. Based on these results, we concluded that loss of the *CjTHI4* gene rendered *C. jadinii* CBS 621 a thiamine auxotroph.

### 6. Cultivation of prototrophic *C. jadinii* strains on ethanol on a vitamin-free medium

To evaluate the physiology of potentially prototrophic *C. jadinii* strains on ethanol, twelve additional strains were selected, excluding those isolated from animal material unless a genome sequence was available. A serial-dilution based method (Fig 4A-B) was used to calculate their specific growth rates in microtiter plates (39). Of the tested *C. jadinii* strains, only CBS 621 and CBS 7232 failed to grow on vitamin-free medium (Fig 4C). Growth of these strains was rescued by adding of thiamine and biotin, respectively (Fig S5A). The maximum specific growth rates of the strains ranged from 0.13 - 0.44 h^-1^ (Fig 4C). Most of the variation in the dataset could be attributed to strain differences (p<0.001). Vitamin absence also significantly impacted the specific growth rates, leading to an average 10-15% lower growth rate (p<0.001) (Fig 4C). The final Green Value of the growth curves (Fig S5B) showed less variation among samples.

**Figure 4:**
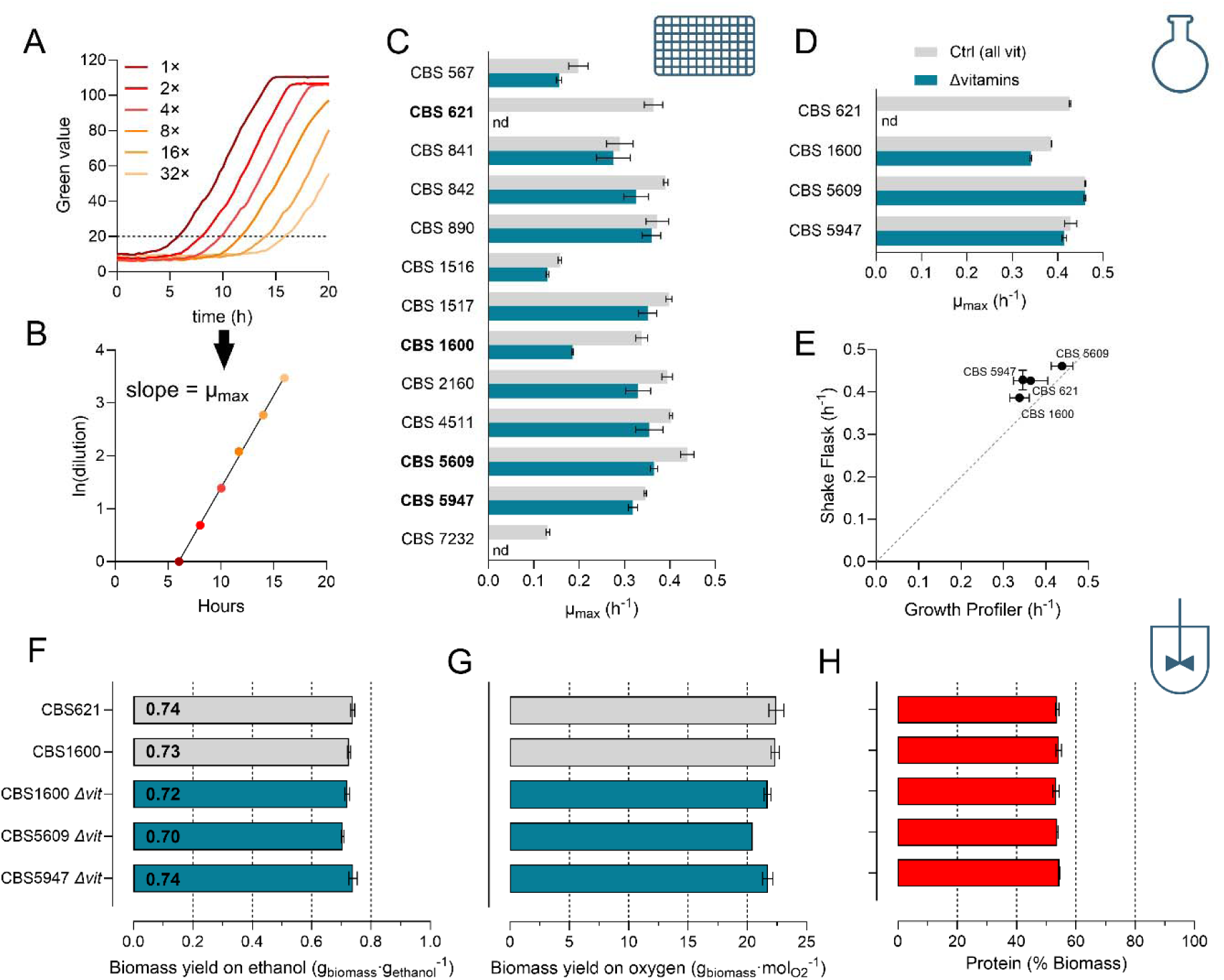
Characterization of *C. jadinii* strains in microtiter plates, shake flasks and bioreactors concerning vitamin dependency. (A) Representative growth curves of serially diluted CBS 621 in microtiter plates, ranging from 1 to 32× dilution, initial OD_660_ 0.2 to 0.00625, respectively. (B) Calculation of specific growth rates via serial dilution method. The time (h) required for a specific dilution to reach a green value of 20 was plotted against the natural logarithm of the respective dilution (1–32). (C) Specific growth rates for 13 *C. jadinii* strains grown on ethanol in the presence or absence of vitamins, calculated with the serial dilution method, n.d: non-detectable. (D) Specific growth rates of 4 *C. jadinii* strains grown on ethanol in shake flasks in the presence or absence of vitamins. (E) Correlation of specific growth rates obtained in microtiter plates (x-axis) with those measured in shake flasks (y-axis) for strains grown in the presence of vitamins. (F-H) Biomass yields on ethanol, oxygen and protein content for selected strains grown in ethanol-limited chemostats (dilution rate 0.10 h^-1^). Where indicated by Δvit, no vitamins were added to the cultivation, n=2-3.

Three *C. jadinii* strains were selected for further investigation: CBS 5609 (highest µ_max_), CBS 1600 (largest difference in µ_max_ caused by vitamin absence) and CBS 5947 (among the top three strains with the highest final Green Value). Maximum specific growth rates of these three strains were measured in shake flasks to validate the serial dilution method. Although the effect of vitamin absence on CBS 1600 was less pronounced (Fig 4D), values obtained with the two methods corresponded well (Fig 4E, S5C). Publicly available genome sequences from strains CBS 1600 and CBS 5609 (24, 79), indicated the presence of a complete *THI4* gene. Presence of *THI4* in strain CBS 5947 was confirmed by whole-genome sequencing (Table S4). Like strains CBS 5609 and CBS 621, *C. jadinii* CBS 5947 appeared to be triploid based on its SNP distribution (Fig S4B).

To investigate biomass yields and protein contents of the three selected strains during growth on ethanol in vitamin-free medium, they were grown in ethanol-limited chemostats at 0.10 h^-1^ in the absence of supplemented vitamins (Table S5). *C. jadinii* CBS 5609, the strain with the highest µ_max_ on ethanol (0.46 h^-1^, Fig 4D) showed the lowest biomass yield on ethanol and oxygen (Fig 4F-G). Strains CBS 1600 and CBS 5947 showed nearly the same biomass yields on ethanol (0.72 and 0.74 g_biomass_·(g_ethanol_)^-1^, respectively (Fig 4F-H). All three strains showed a protein content of approximately 54% (Fig 4H). Vitamin-supplemented ethanol-limited cultures of strains CBS 1600 and CBS 621 showed the same biomass yield on ethanol and protein content (Fig 4F-H). For strain CBS 1600, these results indicate that omitting vitamins from synthetic media does not affect biomass yield and protein content during ethanol-limited growth at 0.10 h^-1^.

### 7. High-cell-density fed-batch cultivation of *C. jadinii* on ethanol in a mineral, vitamin-free medium

Of the vitamin-prototrophic *C. jadinii* strains, CBS 5947 showed the highest biomass yield in ethanol-limited chemostat cultures (0.74 g_biomass_·(g_ethanol_)^-1^, Fig 4F). To evaluate whether this strain could be grown to high biomass densities on ethanol in a vitamin-free medium, a fed-batch protocol was developed. After an initial batch phase, a biomass concentration of 5.5 g_biomass_·(kg_broth_)^-1^ was reached in two independent reactors. For the subsequent fed-batch regime, pure ethanol was continuously fed and 25% NH_4_OH was used as pH titrant and nitrogen source. To increase oxygen solubility, 1 bar overpressure was applied since the beginning of the batch phase (Fig 5A). The initial specific growth rate was set at 0.10 h^-1^ by applying a preprogrammed exponentially increasing ethanol feed rate (Fig 5B). When dissolved oxygen concentration in the reactor decreased to a concentration corresponding to 15% of air saturation at atmospheric pressure (Fig S6A), the ethanol feed rate was first fixed at 0.047 kg_ethanol_·h^-1^ and, after 5 additional hours, at 0.035 kg_ethanol_·h^-1^ (Fig 5B) to prevent oxygen limitation. In total, approximately 1.4 kg of ethanol was consumed in each reactor within 48 h (Fig 5C), leading to a biomass concentration of 98 ± 4 g_biomass_·(kg_broth_)^-1^ (Fig 5D). Consistent with the cultures being ethanol-limited, the ethanol concentration in culture samples remained below detection limit throughout the fed-batch phase. From the time-dependent increase of the amount of biomass in the reactor, we calculated an approximate specific growth rate of 0.02 h^-1^ at the end of the fed-batch phase (Fig 5E). A decline in growth rate was also evident from the slope of the CO_2_ profile during fermentation (Fig S6B). The fed-batch process was simulated (37) based on the parameters calculated from Fig 2D-F and the feed profile shown in Fig 5B. Experimentally determined specific growth rates and ethanol consumption were consistent with model predictions (Fig 5C-E).

**Figure 5:**
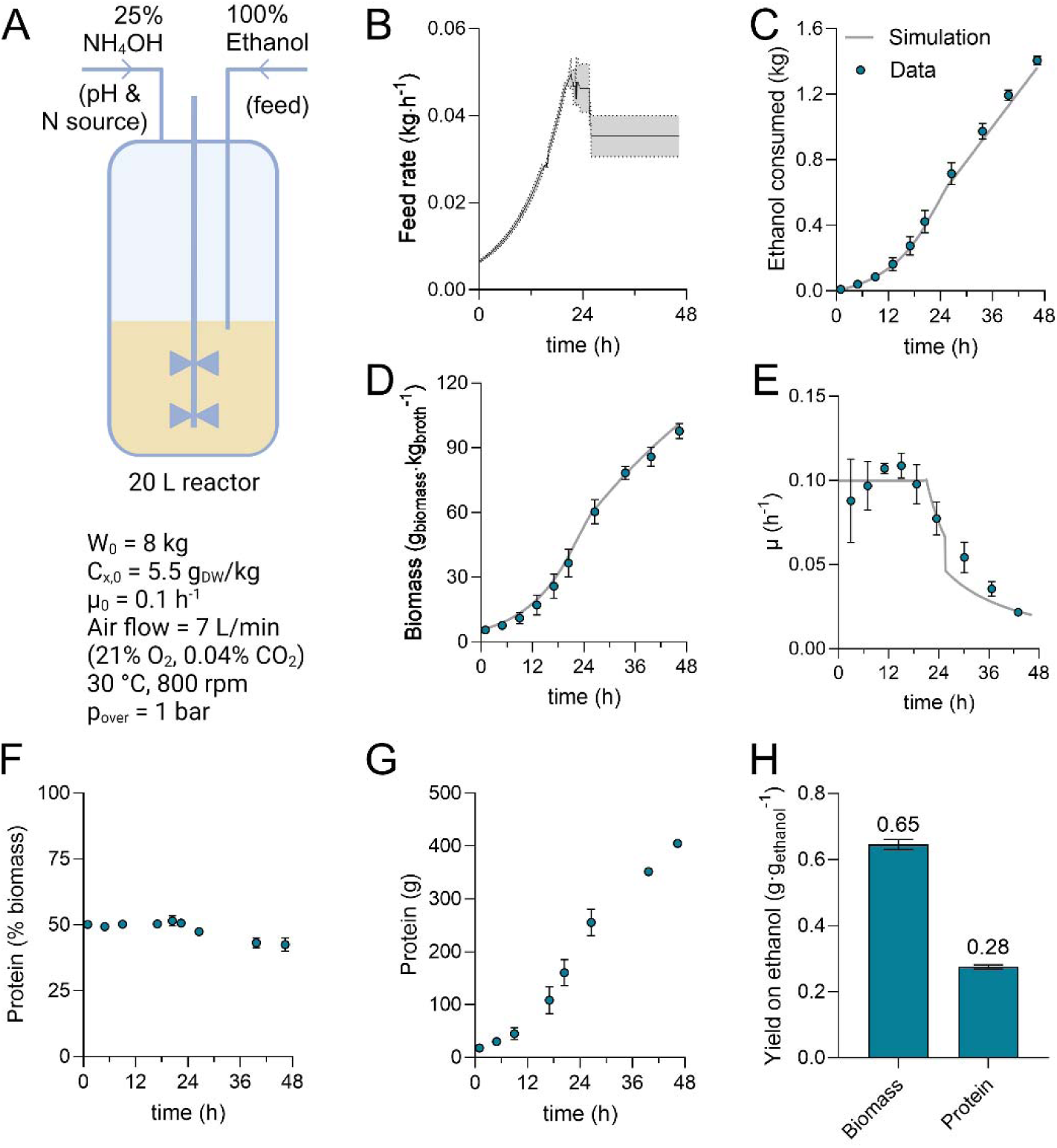
Growth of *C. jadinii* CBS 5947 on vitamin-free medium in aerobic, ethanol-limited fed-batch cultures. Aerobic fed-batch fermentation was conducted at 30 °C, at pH 5.0 and at 1 bar overpressure, with a feed of pure ethanol. A 25% (w/v) NH_4_OH solution was used for simultaneous pH control and nitrogen supplementation. (A) Scheme of reactor and critical parameters related to the process (fed-batch phase). W_0_: initial weight of the culture broth (kg), c_x,0_: initial biomass concentration (kg_biomass_·(kg_broth_)^-1^), µ_0_: initial specific growth rate in fed-batch phase (h^-1^), DW = biomass dry weight concentration (g biomass·(kg_broth_)^-1^). Figure created in BioRender. Vieira-Lara, M. (2024) BioRender.com/z76n618. (B) Profile of feed rate during the fed-batch phase. (C-E) Consumed ethanol, biomass dry weight, and growth rate (µ) during the fed-batch phase. Symbols represent data and grey lines refer to simulations. (F-G) protein percentage in the biomass and total protein content during the fed-batch phase. (H) Total biomass and protein yield on ethanol based on the whole fed-batch process. Data are shown for 2 independent reactors (mean ± SD).

The protein content remained at 50% during the exponential feed phase of the fed-batch culture (Fig 5F). However, in contrast to observations in steady-state chemostat cultures (Fig 2) in which protein concentrations did not decline at low specific growth rates, a decrease of the protein content to 42% was observed towards the end of the fed-batch cultures (Fig 5F). This indicates that the protein content decreased when the growth rate was dynamically changed from 0.10 to 0.022 h^-1^. In the two fed-batch experiments, an average of 405 g protein was produced (Fig 5G). Based on the total ethanol consumed and total biomass and protein produced during the fed-batch phase, biomass, and protein yields of 0.65 g_biomass_·(g_ethanol_)^-1^ and 0.28 g_protein_·(g_ethanol_)^-1^, respectively, were calculated (Fig 5H). Production of low concentrations of acetate and succinate in the fed-batch processes (∼10 mM final concentrations) (Fig S6C) played a negligible role in the total carbon recovery (Fig S6D-E).

In the fed-batch cultures, strain CBS 5947 needed to synthesize vitamins to sustain growth. Levels of thiamine, riboflavin, niacin, and pyridoxine in fed-batch grown *C. jadinii* CBS 5947 ranged from 1.7 – 260 mg ·(kg_biomass_)^-1^ (Table S6). When compared with the vitamin contents in aquaculture diets (80), ethanol-grown *C. jadinii* biomass showed higher levels of niacin, riboflavin and pyridoxine, while thiamine content of the yeast biomass was approximately 3-fold lower (Table S6).

## Discussion

Fast growth of industrial microorganisms is especially beneficial during batch cultivation and during the initial exponential growth phase of aerobic fed-batch fermentation processes, when oxygen transfer is not yet limiting. In a recent screening of 52 ascomycete budding yeasts, we noted that *C. jadinii* CBS 621 was the fastest-growing strains on a synthetic medium containing ethanol as sole carbon source (13). In the present study, multiple *C. jadinii* strains showed specific growth rates on ethanol close to 0.4 h^-1^ (doubling time 1.7 h) in batch cultures grown on a synthetic medium supplemented with a standard vitamin mixture (Fig 2B, 4C-D). With two exceptions, the investigated *C. jadinii* strains were prototrophic and showed maximum specific growth rates on ethanol in vitamin-free media that were only 10-15% lower than in vitamin-supplemented medium. Use of vitamin-free media offers advantages in terms of cost and simplicity of medium preparation and sterilization and, potentially, reduction of contamination risks (78). Moreover, use of a naturally prototrophic yeast strain precludes the need for extensive metabolic engineering strategies, such as recently applied to *S. cerevisiae* (62), to eliminate full vitamin auxotrophies. The thiamine auxotrophy of our initial model strain *C. jadinii* CBS 621 (Fig 3A-C) could be attributed to absence of the *CjTHI4* gene. Thiamine auxotrophy of this strain may enable use of *CjTHI4* as a selectable marker in ongoing research to make *C. jadinii* fully accessible to Cas9-based genome editing (81).

Biomass yields of *C. jadinii* CBS 621 in ethanol-limited chemostat cultures with vitamin supplementation, and of two prototrophic *C. jadinii* strains in vitamin-free media, were among the highest reported for yeasts (13) (Fig 4F). The maximum biomass yield (Y_x/s_^max^) of *C. jadinii* CBS 621 on ethanol of 0.83 g_biomass_·(g_ethanol_)^-1^ in vitamin-supplemented, ethanol-limited chemostat cultures even matched the theoretical maximum yield on ethanol of 0.83 - 0.84 g_biomass_·(g_ethanol_)^-1^ (13, 16) for ethanol-limited growth with ammonium as the nitrogen source. A previous study on strain CBS 621 that was based on measurements at a single dilution rate (0.10 h^-1^) reported lower biomass yields than observed in our study (18). A similarly high biomass yield on ethanol (0.82 g·g^-1^) was reported for batch cultures of the related species *Cyberlindnera saturnus* (12). We consistently measured high biomass yields in replicate cultures of different *C. jadinii* strains, both on vitamin-supplemented and vitamin-free media, checked carbon recoveries of the chemostat cultures and confirmed that no biomass retention occurred (82) and therefore trust our reported data.

Ethanol-limited chemostat cultures of *C. jadinii* showed a 25% higher biomass yield on ethanol than cultures of *S. cerevisiae* grown under the same set of conditions (Fig. 1). This difference can be attributed to the presence and absence, respectively, of genes encoding a proton-pumping Complex I NADH dehydrogenase in these two yeasts (13). In contrast to the difference observed in ethanol-limited cultures (16, 18, 58), biomass yields of *C. jadinii* and *S. cerevisiae* in aerobic glucose-limited cultures differed by only 6% (Fig 1). This observation suggests that the contribution of Complex I to respiratory reoxidation of NADH by *C. jadinii* may be carbon-source dependent. A deeper understanding of the role and regulation of Complex I in *C. jadinii* may contribute to improving yields of biomass and other ATP-intensive products, also on carbon- and energy sources other than ethanol. Our current lack of understanding on this subject is illustrated by the observation that, counterintuitively, levels of the non-proton-pumping single-subunit NADH dehydrogenase *Cj*Ndi1 were approximately 150-fold higher in the proteome of ethanol-limited cultures of *C. jadinii* than in glucose-limited cultures (Fig S1A, Supp Material S1). Further research should also address the question of how subcellular localization of key enzymes and/or redox shuttle mechanisms enable reoxidation of NADH generated by ethanol and acetaldehyde dehydrogenases, enzymes that can occur in the cytosol as well as in the mitochondrial matrix of yeasts (61), by Complex I, whose catalytic site is in the mitochondrial matrix.

The estimated growth-rate independent maintenance energy requirement for ATP (m_ATP_) of ethanol-limited cultures of *C. jadinii* (2.7 mmol ATP·(g_biomass_^-1^)·h^-1^, Fig 2F) was higher than the m_ATP_ of 1.2 mmol ATP·(g_biomass_)^-1^·h^-1^ estimated from a study on aerobic, glucose-limited chemostat cultures of the Complex-I containing yeast *Komagataella phaffii*, grown at dilution rates ranging from 0.025 to 0.10 h^-1^ (83). In energy-substrate-limited cultures, a high maintenance energy requirement most strongly affects biomass yields at low specific growth rates, where it needs a large fraction of the substrate to be respired instead of converted to biomass. Consequently, and consistent with model-based simulations that included the high maintenance requirement of *C. jadinii*, high-cell-density fed-batch cultures did not attain the biomass yields reached in ethanol-limited chemostat cultures grown at intermediate rates (Fig 3C, 4F, 5H). Protein turnover has been reported to account for a large share of the ATP requirement for cellular maintenance in *S. cerevisiae* (84). Interestingly, two heterologous-protein producing *K. phaffii* strains were reported to exhibit 2-fold higher maintenance-energy requirements than a reference strain (83, 85). This raises the question of whether the high m_ATP_ of *C. jadinii* may be related to its high protein content, and particularly the remarkably high levels of specific proteins involved in ethanol metabolism.

Under substrate-limited conditions, competitiveness of microorganisms is determined by their affinity for the growth-limiting substrate, i.e. the ability to sustain a high biomass-specific consumption rate (q_s_) at a low concentration (c_s_) of that substrate. Based on the Monod equation (q_s_ = q_s,max_ (c_s_/(c_s_ + K_s_))), affinity can be defined as q_s,max_/K_S_ (86), in which q_s,max_ is the maximum biomass-specific substrate uptake rate, and K_s_ is the value of c_s_ at which q_s_ = ½·q_s,max_. At low specific growth rates, a large fraction of the proteome of ethanol-limited cultures of *C. jadinii* was allocated to enzymes that catalyse the first three reactions of ethanol metabolism (Fig 2I) and, in particular, to alcohol dehydrogenase (Fig 2J). This observation indicates that, at low specific growth rates, *C. jadinii* increases its affinity for ethanol by increasing q_s,max_. This adaptation strategy resembles regulation of methanol oxidase, the first enzyme in methanol metabolism, in *Ogataea polymorpha*. In slow-growing (0.03 h^-1^) methanol-limited cultures of this methylotrophic yeast, 20% of its proteome consisted of methanol oxidase (87), which closely resembles the 17% contribution of alcohol-dehydrogenases in ethanol-limited *C. jadinii* cultures grown at 0.025 h^-1^. We hypothesize that this ‘q_s,max_ adaptation’ to ethanol-limited growth contributes to the observation that, despite lower rates of assimilation and dissimilation at low specific growth rates, *C. jadinii* exhibited an almost growth-rate independent protein content of ca. 54% in ethanol-limited chemostat cultures.

Based on the quantitative data obtained from chemostat experiments, we designed a feed profile for high-cell density ethanol-limited fed-batch cultures. A previous study also described based fed-batch cultivation of *C. jadinii* on ethanol, but applied discontinuous ethanol feeding and reached a maximum biomass concentration of 15 g_biomass_·(kg_broth_)^-1^ (88). By controlled feeding of pure ethanol, which eliminated the need for sterilisation of the feed line, the fed-batch cultures reached 100 g_biomass_·(kg_broth_)^-1^ in 48 h of cultivation (Fig 5D). Experimental data on biomass formation were in excellent agreement with model predictions. However, the protein content of biomass harvested at the end of high-cell density fed-batch cultures, when the specific growth rate had decreased to 0.02 h^-1^, had decreased to only 42%. Further research is needed to assess whether this marked difference with the protein content in slow-growing chemostat cultures is due to the dynamic growth-rate profile in the fed-batch cultures, imperfect mixing of the pure ethanol feed, or other factors.

## Conclusions

Based on different analyses on ethanol-grown batch, chemostat and fed-batch cultures, the present study provides a quantitative physiology baseline for fundamental and applied research on *C. jadinii*. Its high maximum growth rate, high energetic efficiency of ethanol dissimilation, simple nutritional requirements and high protein content, make this yeast a highly interesting platform for production of SCP and other products from ethanol produced by low-emission technologies. Especially for SCP applications, an observed difference in protein content between ethanol chemostat and fed-batch cultures that were grown over the same range of specific growth rates, merits further investigation. From a fundamental science perspective, our results identify questions about regulation, energy coupling and compartmentation of NADH reoxidation during growth of yeasts and fungi during growth on non-conventional substrates. In addition, the results underline the key role of maintenance energy requirements in high-cell-density fed-batch processes.

## Declarations

### Ethics approval and consent to participate

Not applicable

### Consent for publication

Not applicable

### Availability of data and materials

All data generated or analysed during this study are included in this published article and its supplementary information files.

### Authors’ contributions

MAVL: investigation, methodology, analysis, data visualization, writing (original draft, reviewing and editing). MW: investigation, methodology, analysis, writing (reviewing and editing). EdH and MvdB: investigation, methodology. JMD: conceptualization, writing (reviewing and editing). JTP: conceptualization, supervision, writing (reviewing and editing).

## Acknowledgements

The authors thank Clara Carqueija Cardoso for the help with DNA sequencing, Christiaan Mooiman for assistance with the fed-batch experiments, Marijke Luttik for the help with growth experiments, Chris Klomp and Niels Tiemersma for evaluating the methods for biomass composition and help with bioreactor cultivation, and Jordy Elfering for the testing of the serial dilution method and screening and for help with bioreactor cultivation. We thank André Vente for the amino acid and proteome analyses, and Mickel Jansen and Tjeerd van Rij for the input in the project.

