## Supplemental Materials for "Quantitative physiology and biomass composition of *Cyberlindnera jadinii* in ethanol-grown cultures"

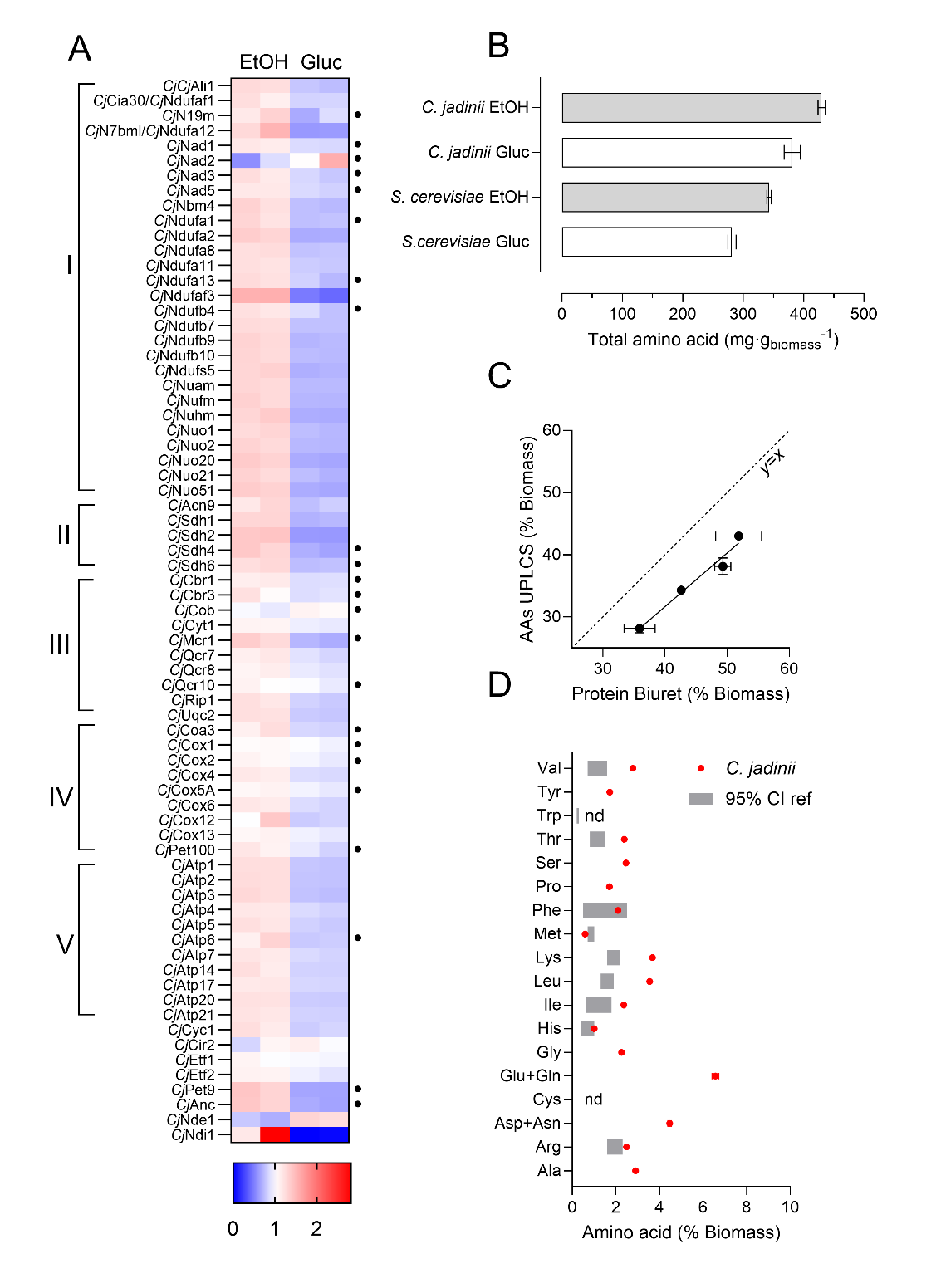

**Figure S1: Amino acid composition and OXPHOS proteins.** (A) Heatmap of proteins involved in oxidative phosphorylation (OXPHOS) in ethanol- (EtOH) vs glucose-grown (Gluc) *C. jadinii*. Levels were normalized to average levels across all samples. Blue: low levels, red: high levels. Protein names and UniProt ID’s are found in Supp. Material S1. Dots next to each row on the right represent predicted membrane proteins. Roman numerals on the left represent respiratory complex (I-IV) and F_1_F_o_-ATP synthase (V). (B) Total amino acid residue concentration in *S. cerevisiae* and *C. jadinii* grown on glucose or ethanol as substrate in chemostats. (C) Comparison of total amino acid levels (y-axis) and total protein measured with the Biuret method (x-axis). The dashed line is the identity line (y=x). (D) Amino acid levels in ethanol-grown *C. jadinii* (red dots) compared to the 95% confidence interval (CI) of amino acid levels needed for feed formulation in aquaculture (from (72)), grey boxes, n =2.

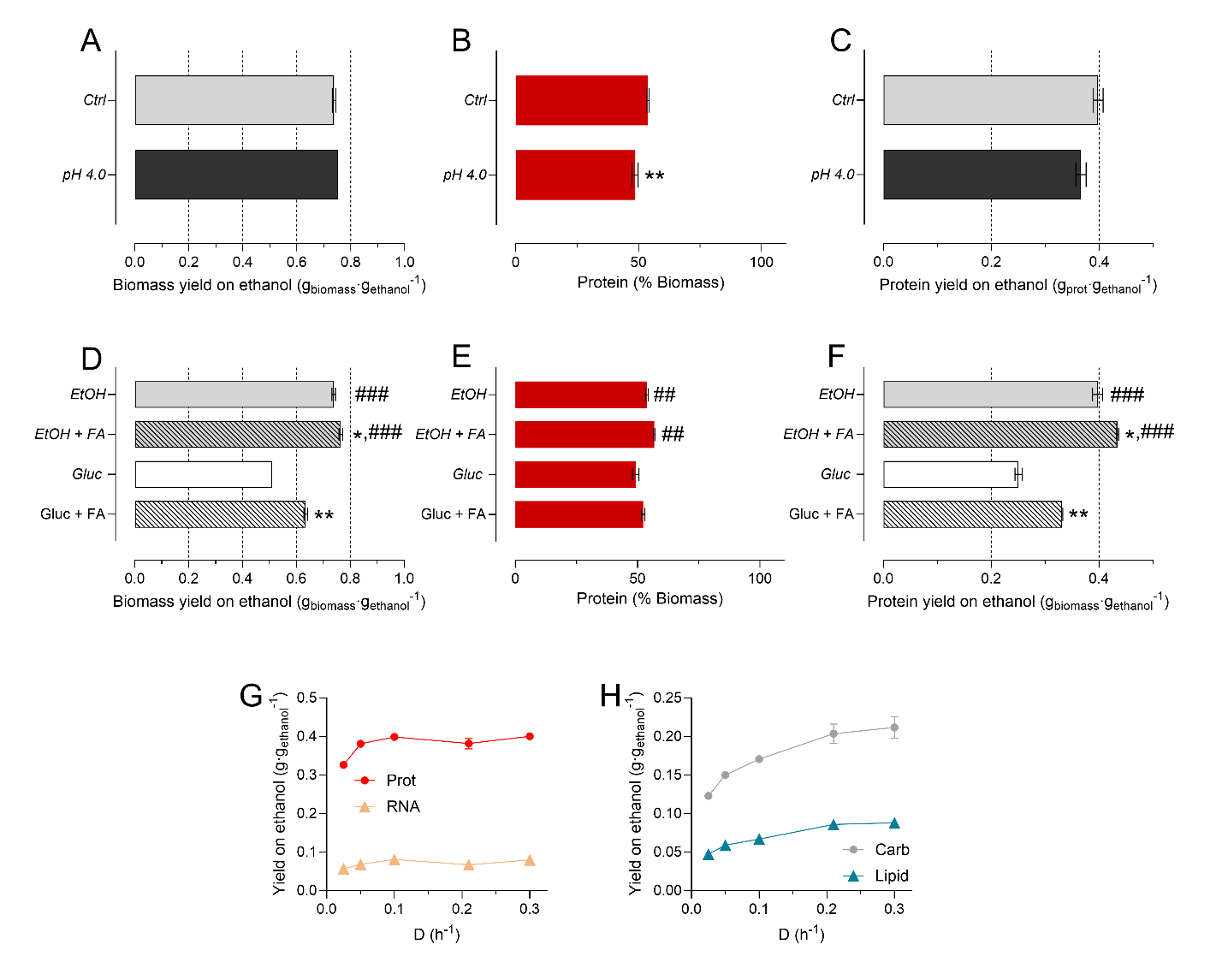

**Figure S2: Yields of biomass components and strategies to increase biomass yield** (A) Biomass yield on substrate, (B) protein content as a percentage of the biomass and (C) protein yields on substrate for *C. jadinii* grown in ethanol-limited cultures at either pH 5.0 (Ctrl) or pH 4.0. (D) Biomass yield on substrate, (E) protein content as a percentage of the biomass and (F) protein yields on substrate for *C. jadinii* grown in ethanol (EtOH) or glucose-limited (Gluc) chemostats in the presence or absence of formic acid (FA). The ratio of formic acid to substrate was defined as 1.2 Cmol/Cmol: 2.4 mol FA per mol ethanol and 7.2 mol FA per mol glucose. (G) and (H) Yields of the main components of the biomass in g∙g_s_^‑1^: proteins (Prot), RNA, carbohydrates (Carb) and lipids, ##p<0.01 ###p<0.001, ethanol- (EtOH) vs glucose-grown (Gluc). *p<0.05, **p<0.01, effect of either pH or FA addition, n=2.

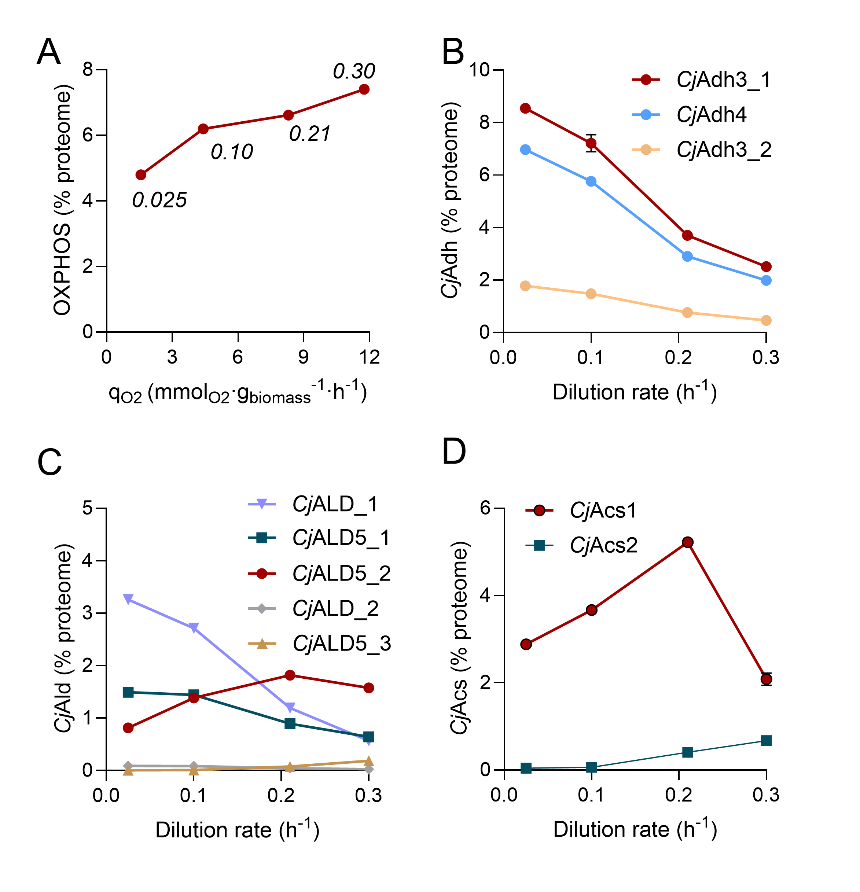

**Figure S3: Proteome allocation of OXPHOS and ethanol catabolism proteins.** (A) Percentage of the proteome occupied by OXPHOS proteins versus the oxygen consumption rates (q_O2_). Indicated in italics are the respective dilution rates. (B) Alcohol dehydrogenase (*Cj*Adh) isoforms, (B) aldehyde dehydrogenase isoforms (*Cj*Ald) and (C) acetyl-CoA synthetase isoforms (*Cj*Acs) as a function of the dilution rate. Protein names and UniProt ID’s are found in Supp. Material S2, n =2.

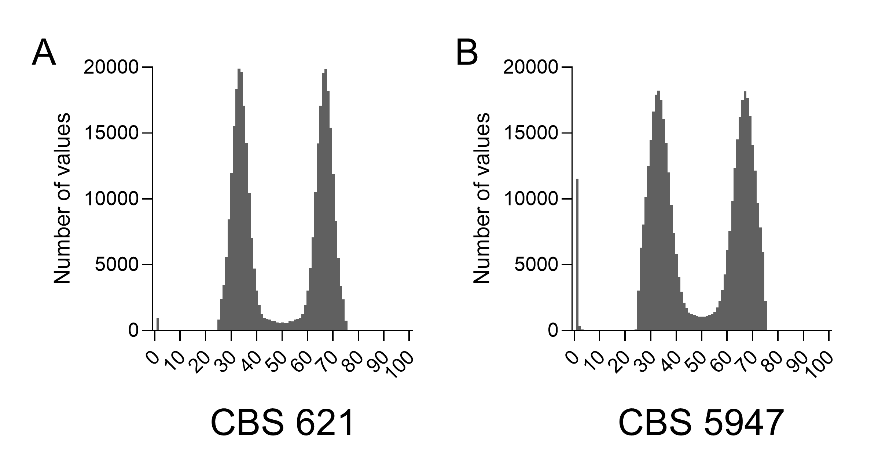

**Figure S4:** SNP distribution. Reads were mapped to the haploid consensus genome. SNPs were detected and the percentage of reads (vs total reads) for each SNP were computed and plotted as histograms. The peaks at around 33% and 66% show that SNPs occupy either 1/3 or 2/3 of each location, indicating that (A) CBS 621 and (B) CBS 5947 are triploid strains.

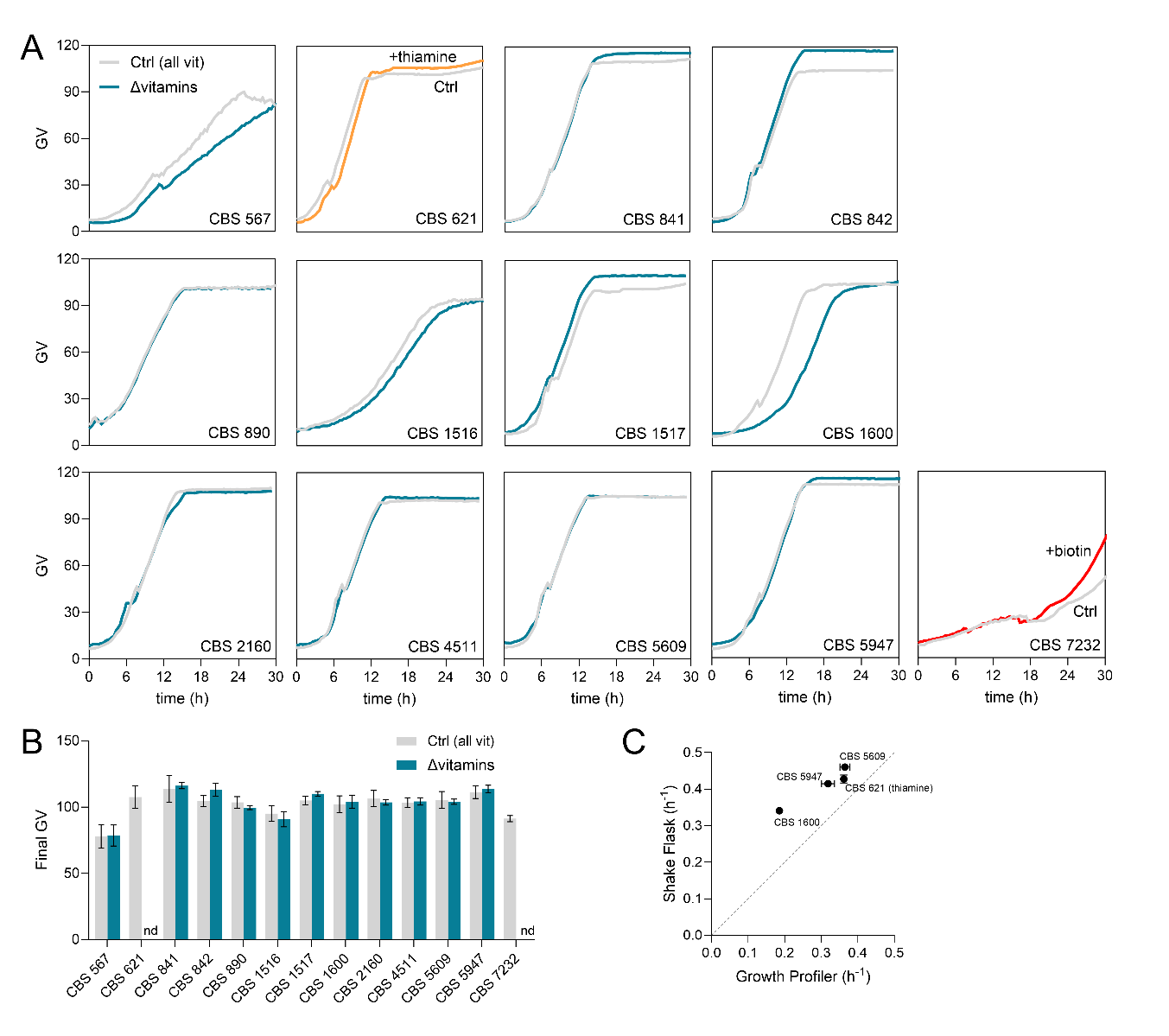

**Figure S5: Growth curves of 13 *C. jadinii* strains.** (A) Representative traces of individual growth curves for all tested strains. Traces represent cultures grown at dilution 1×, OD_initial_ = 0.2. Grey lines represent the control (with added vitamins), blue lines represent cultures grown in the absence of added vitamins. For CBS 621 and CBS 7232, which do not grow in the absence of all vitamins, thiamine and biotin were added as sole vitamins, respectively. (B) Final green values (GV) at the stationary phase for all conditions, nd: non-detectable. (C) Correlation of the growth rates obtained in the Growth Profiler (x-axis) with those obtained in shake flasks (y-axis) for strains grown in the absence of vitamins, n=2-3.

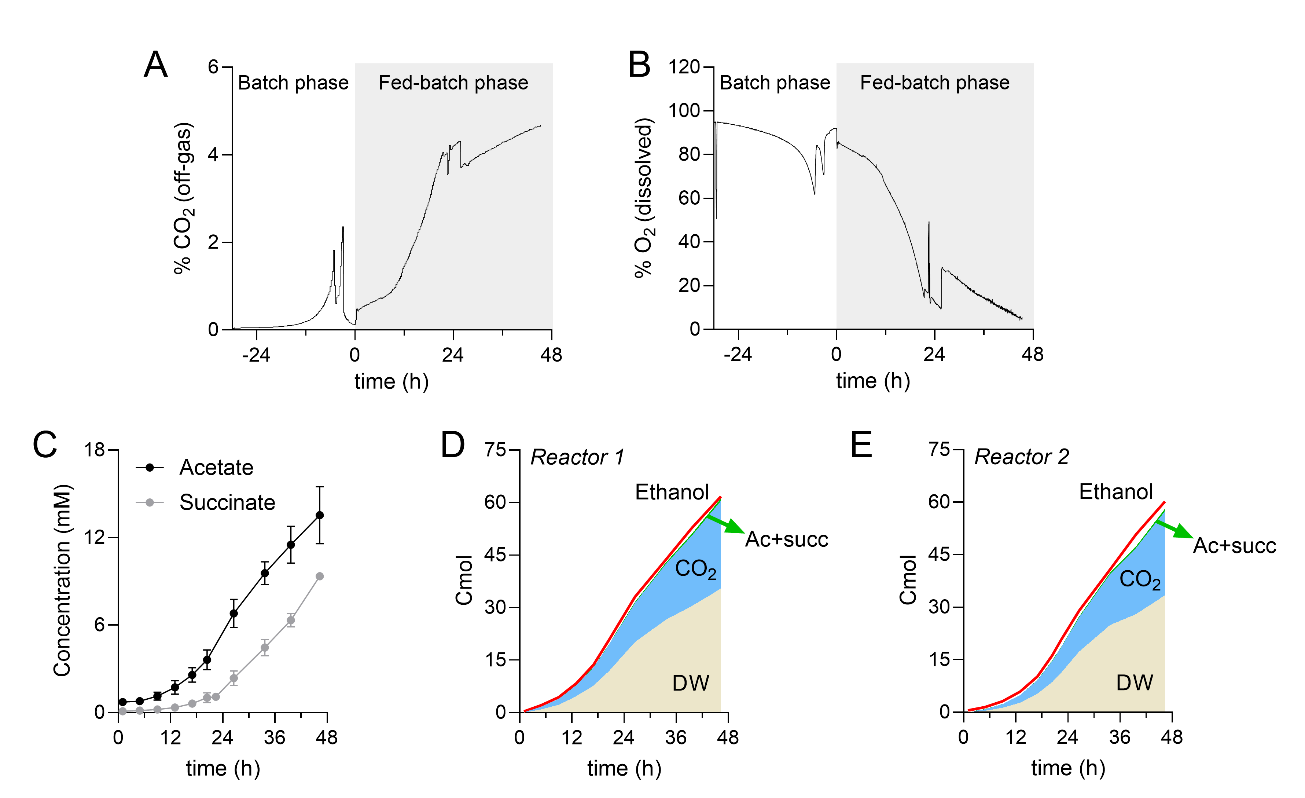

**Figure S6: Fed-batch representative traces and carbon recovery.** (A) Representative CO_2_ concentration in the off-gas and (B) dissolved oxygen (DO) concentration, both during batch and fed-batch phases for reactor 2. (C) Acetate and succinate concentrations in the residuals during the fed-batch phase. (D-E) carbon balance time course for reactors 1 and 2, respectively. The red line represents the total amount of ethanol consumed. The shaded areas correspond to biomass (DW), CO_2_ and organic acids (Ac + succ). All amounts are expressed in Cmol, n=2.

| Strain | Substrate | Y_X/S_  (g_biomass_∙g_s_^-1^) | q_ethanol_  (mmol∙g_biomass_^-1^∙h^-1^) | q_CO2_  (mmol∙g_biomass_^-1^∙h^-1^) | q_O2_  (mmol∙g_biomass_^-1^∙h^-1^) | RQ |
| --- | --- | --- | --- | --- | --- | --- |
| *S. cerevisiae* | Glucose | 0.48 ± 0.00 | 1.27 ± 0.00 | 3.31 ± 0.25 | 2.78 ± 0.13 | 1.19 ± 0.03 |
| *S. cerevisiae* | Ethanol | 0.59 ± 0.01 | 3.64 ± 0.20 | 3.56 ± 0.10 | 6.62 ± 0.17 | 0.54 ± 0.00 |
| *C. jadinii* | Glucose | 0.51 ± 0.01 | 1.08 ± 0.01 | 2.99 ± 0.08 | 2.51 ± 0.12 | 1.19 ± 0.03 |
| *C. jadinii* | Ethanol | 0.74 ± 0.01 | 2.87 ± 0.02 | 2.19 ± 0.02 | 4.41 ± 0.05 | 0.50 ± 0.00 |

**Supplemental Tables**

**Table S1: *C. jadinii* and *S. cerevisiae* in glucose- and ethanol-limited growth at 0.10 h^-1^ (mean ± SD)**

|  | *S. cerevisiae* | | | *C. jadinii* | | |
| --- | --- | --- | --- | --- | --- | --- |
| Amino acid | | **Glucose** | **Ethanol** | | **Glucose** | **Ethanol** |
| *Alanine* | | 17.7 ± 0.5 | 21.4 ± 0.3 | | 25.0 ± 0.8 | 29.1 ± 0.4 |
| *Arginine* | | 15.5 ± 0.4 | 19.5 ± 0.3 | | 21.2 ± 0.6 | 24.9 ± 0.3 |
| *Aspartate/Asparagine* | | 32.9 ± 0.8 | 37.8 ± 0.1 | | 40.5 ± 1.3 | 44.7 ± 0.8 |
| *Glutamate/Glutamine* | | 38.2 ± 0.9 | 60.5 ± 0.6 | | 62.0 ± 2.8 | 65.7 ± 1.6 |
| *Glycine* | | 13.8 ± 0.3 | 16.7 ± 0.0 | | 19.6 ± 0.6 | 22.6 ± 0.1 |
| *Histidine* | | 6.5 ± 0.2 | 8.1 ± 0.1 | | 8.2 ± 0.3 | 10.0 ± 0.1 |
| *Isoleucine* | | 15.8 ± 0.4 | 18.1 ± 0.1 | | 20.4 ± 0.7 | 23.6 ± 0.1 |
| *Leucine* | | 24.0 ± 0.6 | 27.8 ± 0.4 | | 31.4 ± 1.0 | 35.5 ± 0.4 |
| *Lysine* | | 25.8 ± 0.6 | 28.7 ± 0.3 | | 31.9 ± 1.2 | 36.8 ± 0.4 |
| *Methionine* | | 5.1 ± 0.1 | 5.9 ± 0.2 | | 5.2 ± 0.2 | 5.9 ± 0.1 |
| *Phenylalanine* | | 14.5 ± 0.4 | 16.4 ± 0.2 | | 18.8 ± 0.6 | 21.0 ± 0.2 |
| *Proline* | | 11.8 ± 0.3 | 13.8 ± 0.0 | | 15.0 ± 0.4 | 17.1 ± 0.4 |
| *Serine* | | 15.7 ± 0.4 | 17.9 ± 0.4 | | 22.2 ± 1.0 | 24.6 ± 0.4 |
| *Threonine* | | 16.0 ± 0.4 | 18.2 ± 0.2 | | 21.7 ± 0.8 | 23.9 ± 0.4 |
| *Tyrosine* | | 11.0 ± 0.3 | 12.7 ± 0.1 | | 14.9 ± 0.2 | 17.2 ± 0.3 |
| *Valine* | | 17.5 ± 0.5 | 19.6 ± 0.3 | | 23.9 ± 0.8 | 27.8 ± 0.2 |

**Table S2: Amino acid composition of the biomass of *C. jadinii* and *S. cerevisiae* in glucose- and ethanol-limited growth at 0.10 h^-1^ (mean ± SD). Values are in mg∙g_biomass_^-1^.**

| Dilution rate (h^-1^) | Y_X/S_  (g_biomass_∙g_ethanol_^-1^) | q_ethanol_  (mmol∙g_biomass_^-1^∙h^-1^) | q_CO2_  (mmol∙g_biomass_^-1^∙h^-1^) | q_O2_  (mmol∙g_biomass_^-1^∙h^-1^) | RQ |
| --- | --- | --- | --- | --- | --- |
| 0.025 ± 0.00 | 0.59 ± 0.00 | 0.93 ± 0.02 | 0.98 ± 0.03 | 1.56 ± 0.00 | 0.63 ± 0.02 |
| 0.05 ± 0.00 | 0.70 ± 0.01 | 1.54 ± 0.00 | 1.39 ± 0.01 | 2.65 ± 0.11 | 0.52 ± 0.00 |
| 0.10 ± 0.00 | 0.74 ± 0.01 | 2.87 ± 0.02 | 2.19 ± 0.02 | 4.41 ± 0.05 | 0.50 ± 0.00 |
| 0.21 ± 0.00 | 0.77 ± 0.01 | 5.90 ± 0.01 | 3.63 ± 0.02 | 8.30 ± 0.02 | 0.44 ± 0.00 |
| 0.24 ± 0.00 | 0.78 ± 0.01 | 6.69 ± 0.01 | 3.86 ± 0.04 | 9.72 ± 0.25 | 0.40 ± 0.01 |
| 0.30 ± 0.00 | 0.81 ± 0.00 | 8.18 ± 0.08 | 4.78 ± 0.11 | 11.76 ± 0.07 | 0.41 ± 0.00 |

**Table S3: effect of growth rate on ethanol-limited CBS 621 physiology**

**Table S4: Genome sequence of CBS 621 and CBS 5947**

| Components | NBRC 0988 (CBS 5609) | NBRC 0988 (CBS 5609) | CBS 621 | CBS 5947 |
| --- | --- | --- | --- | --- |
| NCBI assembly reference | GCA_000328385.1 (2016) | GCA_024346625.1 (2022) |  |  |
| Assembly level | Chromosomes | Chromosomes | - | - |
| Genome size | 14.3 Mb | 13.1 Mb | 17.9 Mb | 14.7 Mb |
| Genes | 8,864 | - | 10,949 |  |
| No of scaffolds | 1,002 | 7 | - | - |
| Scaffolds N50 | 189,765 | 2,424,584 | - | - |
| No of contigs | 1,163 | 7 | 7,832 | 6,831 |
| Contigs N50 | 158,681 | 2,424,584 | 3,994 | 3,299 |
| No of chromosomes | 13 | 6 (+ mtDNA) | - | - |
| GC-content | 44.7 | 44.5 | 44.5 | 44.3 |
| Total of CDS | 8,646 | - |  |  |

**Table S5: Vitamin-dependent physiology of *C. jadinii* strains on ethanol-limited growth at 0.10 h^-1^**

| Strain | Condition | Y_X/S_  (g_biomass_∙g_ethanol_^-1^) | q_ethanol_  (mmol∙g_biomass_^-1^∙h^-1^) | q_CO2_  (mmol∙g_biomass_^-1^∙h^-1^) | q_O2_  (mmol∙g_biomass_^-1^∙h^-1^) | RQ |
| --- | --- | --- | --- | --- | --- | --- |
| CBS 621 | Ctrl (all vit) | 0.74 ± 0.01 | 2.90 ± 0.02 | 2.19 ± 0.02 | 4.40 ± 0.05 | 0.50 ± 0.00 |
| CBS 1600 | Ctrl (all vit) | 0.73 ± 0.01 | 2.94 ± 0.05 | 2.19 ± 0.04 | 4.41 ± 0.12 | 0.50 ± 0.00 |
| CBS 1600 | ∆ vitamins | 0.72 ± 0.01 | 2.96 ± 0.00 | 2.25 ± 0.01 | 4.51 ± 0.01 | 0.50 ± 0.00 |
| CBS 5609 | ∆ vitamins | 0.70 ± 0.00 | 3.14 ± 0.16 | 2.42 ± 0.10 | 4.99 ± 0.23 | 0.48 ± 0.00 |
| CBS 5947 | ∆ vitamins | 0.74 ± 0.01 | 2.95 ± 0.03 | 2.23 ± 0.05 | 4.61 ± 0.09 | 0.48 ± 0.00 |

**Table S6: Vitamin levels in C. jadinii CBS 5947 at the end of fed-batch and their requirements in aquaculture (DOI: 10.17226/13039)**

| Vitamin | Content*  (mg∙(kg _biomass_)^-1^) | | Aquaculture#  (mg∙(kg_diet_)^-1^) |
| --- | --- | --- | --- |
| Thiamine | 1.7 ± 0.3 | 5 ± 2 | |
| Riboflavin | 15 ± 3 | 7 ± 1 | |
| Niacin | 290 ± 140 | 30 ± 15 | |
| Pyridoxine | 53 ± 9 | 7 ± 1 | |

*mean ± SD, ^#^mean ± SEM

**Supplemental Materials**

**Supplemental Material S1: Proteomics dataset *S. cerevisiae* and *C. jadinii* (xlsx)**

**Supplemental Material S2: Proteomics dataset *C. jadinii* dilution rates (xlsx)**

**Supplemental Material S3: Derivation of growth rates from serial dilution method**

The Growth Profiler is an equipment used for high throughput screening for growth of strains and medium conditions. However, it produces Green Values (GV), which are based on pictures taken from the bottom of the wells. The value of GV is proportional to OD, but not linearly (Fig. S7). To avoid the need of creating an GV to OD calibration curve for each strain tested, we propose a method to derive maximal growth rates (µ_max_­) based on serial sample dilutions, inspired by the method described by (35).

The method is based on serial dilutions of a culture sample to determine its µmax­ in given conditions. First, a GV in the early exponential phase for each growth curve is selected, namely GV_select_. In the current manuscript, GV_select_ of 20 was chosen, which will always refer to the same OD for the same species and conditions, given that OD is a function of GV (Fig S7). The time required to reach this specific GV_select_ is recorded for each dilution (t_dilution_). By plotting t_dilution_ (x-axis) against the natural logarithm of the dilution factor (y-axis), one can obtain the µ_max_ as the slope of the fitted line. The mathematical reasoning behind this is addressed below.

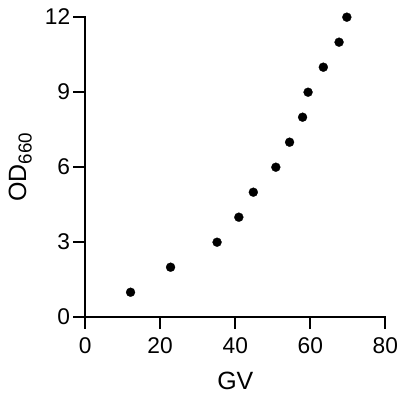

**Figure S7: GV to OD conversion curve, obtained for exponentially-growing CBS 621.**

The general formula for exponential growth at constant volume is:

$c_{t}=c_{0}\cdot e^{\mu\cdot t}$ (S1)

In which c_t_ is the cell concentration at elapsed time, c_0_ is the starting concentration, µ is the specific growth rate, and t is the elapsed time. Let us consider two cultures of the same species, one inoculated at concentration c_n_ and other inoculated at concentration c_p_, where n and p are the dilution factors in respect to c_0_, n>p. After a timespan of t_n_ and t_p_­, respectively, they reach a predetermined concentration c_t_. Therefore, we can write:

$c_{n}\cdot e^{\mu\cdot t_{n}}=c_{p}\cdot e^{\mu\cdot t_{p}}$ (S2)

$c_{p}=c_{n}\cdot\frac{n}{p}$ (S3)

By substituting c_p_ in equation S2 according to S3, we have:

$e^{\mu\cdot t_{n}}=\frac{n}{p}\cdot e^{\mu\cdot t_{p}}$ (S4)

Solving this equation for µ, we have:

$\mu=\frac{\ln n-\ln p}{\left( t_{n}- t_{p} \right)}$ (S5)

If cells are growing at a µ = µ_max_­, this can then be rewritten as:

$\mu_{max}=\frac{\ln n-\ln p}{\left( t_{n}- t_{p} \right)}$ (S6)

Fig. S8A represents the growth curves (in GV) of a specific strain/condition inoculated at dilutions n and p and the GV_select_. Fig S8B shows a line where the natural logarithms of dilutions (y-axis) are plotted against the time to reach GV_select_ (x-axis). The slope of the curve is equal to equation S6, therefore justifying the validity of the serial dilution method to infer growth rates. This method, however, assumes that no lag phase is present when diluting the samples.

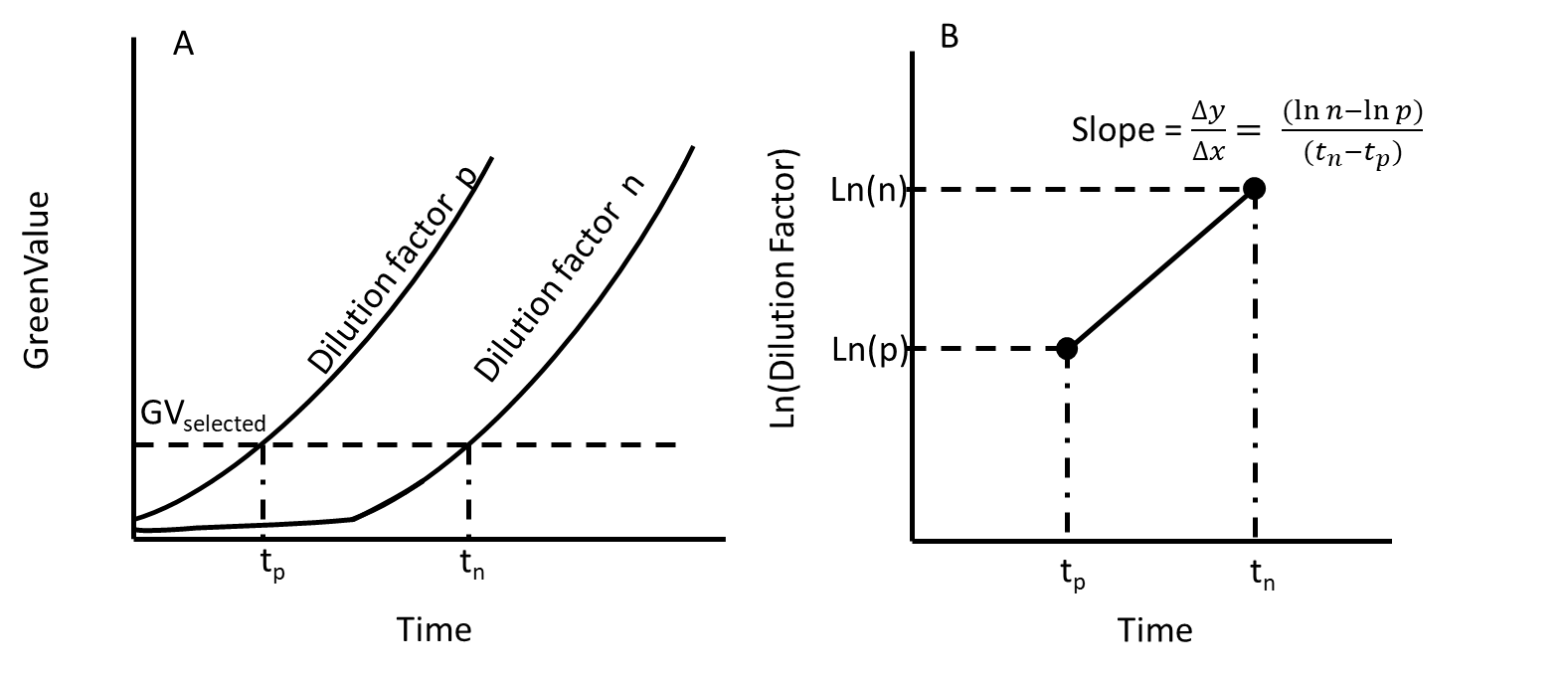

**Fig S8: Schematic graphs of the serial dilution method.** (A) Growth curves of cultures inoculated at dilution factors p and n, in Green Value against time. Time points t_p_ and t_n_ stand for the time when GV_select_ is reached. (B) Natural logarithm of dilution factors p and n plotted against the respective time points t_p_ and t_n_, displayed with the mathematical formula for the slope.
